# Decoding Pain: Uncovering the Factors that Affect Performance of Neuroimaging-Based Pain Models

**DOI:** 10.1101/2023.12.22.573021

**Authors:** Dong Hee Lee, Sungwoo Lee, Choong-Wan Woo

## Abstract

Neuroimaging-based pain biomarkers, when combined with machine learning techniques, have demonstrated potential in decoding pain intensity and diagnosing clinical pain conditions. However, a systematic evaluation of how different modeling options affect model performance remains unexplored. This study presents results from a comprehensive literature survey and benchmarking analysis. We conducted a survey of 57 previously published articles that included neuroimaging-based predictive modeling of pain, comparing classification and prediction performance based on the following modeling variables—the levels of data, spatial scales, idiographic vs. population models, and sample sizes. The findings revealed a preference for population-level modeling with brain-wide features, aligning with the goal of clinical translation of neuroimaging biomarkers. However, a systematic evaluation of the influence of different modeling options was hindered by a limited number of independent test results. This prompted us to conduct benchmarking analyses using a locally collected functional Magnetic Resonance Imaging (fMRI) dataset (*N* = 124) involving an experimental thermal pain task. The results clearly demonstrated that data levels, spatial scales, and sample sizes significantly impact model performance. Specifically, incorporating more pain-related brain regions, increasing sample sizes, and using less data averaging in training while increasing it in testing enhanced performance. These findings provide a useful reference for decision-making in the development of neuroimaging-based biomarkers, highlighting the importance of the careful selection of modeling strategies and options to build better-performing neuroimaging pain biomarkers. These findings offer useful guidance for developing neuroimaging-based biomarkers, underscoring the importance of strategic selection of modeling approaches to build better-performing neuroimaging pain biomarkers.

## Introduction

Neuroimaging-based biomarkers of pain are increasingly gaining attention in basic and clinical studies of pain (Davis et al., 2017; Tracey, 2017). In clinical settings, pain assessment largely depends on self-report, such as the numeric rating scale (Williamson & Hoggart, 2005), but self-report has limitations in the accuracy and clinical utility because pain is a complex biopsychosocial phenomenon that requires multiple types of data for accurate evaluation (Tracey et al., 2019). Among many data types, neuroimaging data provide a unique window that allows us to assess pain based on brain structure and functions, having the potential to serve as biomarkers for the prediction of pain intensity and the diagnosis of clinical pain conditions. Previous studies have compared the classification and prediction performances of neuroimaging-based biomarkers (Woo, Chang, et al., 2017), with one paper focusing on pain (van der Miesen et al., 2019). These studies, however, simply conducted the comparisons without considering the potential impacts of different modeling options and targets on the model performances. In this study, we address the question of how modeling targets and options influence the prediction performances of neuroimaging pain biomarkers through a systematic literature survey and benchmarking analyses.

Figure 1 illustrates the workflows of the literature survey and benchmarking analysis. We compared model performances focusing on various modeling targets (e.g., pain versus no pain classification, pain rating prediction, etc.), data levels (e.g., trial-level, condition-level, etc.), spatial scale (single region, multiple regions, whole-brain, etc.), model levels (e.g., idiographic, population-level, etc.), and sample sizes. These variables were chosen based on their significance in prior research. For example, due to a high level of noise in neuroimaging data, researchers usually average data across multiple trial repetitions to enhance the signal-to-noise ratio. However, it is unclear whether data averaging is always beneficial to model performance. While machine learning algorithms generally necessitate substantial data to recognize meaningful patterns hidden in the data, averaging decreases the quantity of data, potentially along with informative variances, in favor of an improved signal-to-noise ratio. Previous studies commonly suggested that increased data averaging improves classification accuracy and explained variance in predictive models (Lindquist et al., 2017; Woo, Schmidt, et al., 2017). However, these studies typically averaged both training and testing data in a similar manner to ensure consistent data distribution across training and testing sets. Consequently, the impact of data averaging on the model generalizability, particularly when the test data have a different data distribution, requires further investigation.

**Figure 1.**
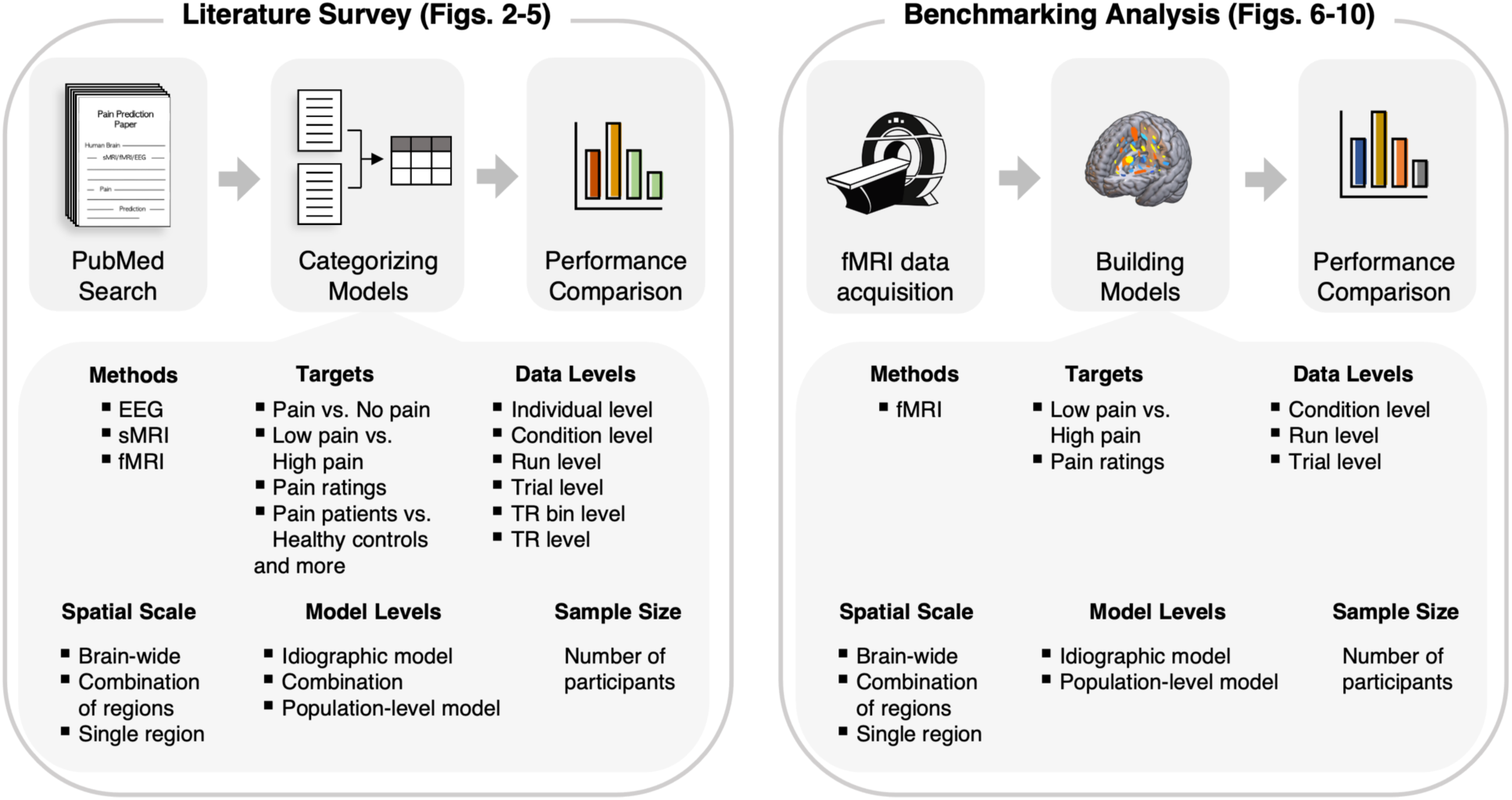
Study Overview. The current study used two approaches to investigate neuroimaging-based pain biomarkers: (1) literature survey and (2) benchmarking analysis. For literature survey, we conducted a literature search on neuroimaging-based biomarkers of pain on PubMed. We then categorized and summarized the predictive models of pain in terms of 12 aspects, including neuroimaging methods, modeling targets, levels of data, spatial scale, levels of model, their sample sizes, tasks, algorithms, etc. In this figure, we displayed more details of 6 aspects among them. Each aspect contained multiple categories. We then compared model performances based on the selected aspects. For benchmarking analysis, we analyzed a large-scale task-based fMRI data with painful thermal stimulation. We developed multiple predictive models with varying modeling options that were appeared important in the literature survey and evaluated the influences of these modeling options on model performance.

Furthermore, it is well-known that the brain representations of pain are distributed across multiple brain systems and processed through a degenerate mechanism (Coghill, 2020; Coghill et al., 1999). Consistent with this idea, previous studies have shown that predictive models containing more voxels and regions show better model performance (Brodersen et al., 2012; Kragel et al., 2018; Petre et al., 2022). However, it is unclear whether including more voxels and regions is always beneficial to model performance, given that it will also increase the possibility of overfitting and introducing more noise. Therefore, we need to conduct a more fine-grained investigation of to what extent the inclusion of voxels and regions will help increase model performance. In addition, it is a common assumption that idiographic models (i.e., individualized predictive models) would better perform in capturing pain ratings of each individual compared to population-level models. However, it is unclear how much data are required to capture the within-individual variability enough to ensure the generalizability of idiographic models. With a mediocre amount of data, it is possible that idiographic models do not outperform population-level models. Lastly, studies have been suggesting larger sample sizes benefit predictive modeling (Marek et al., 2022), but researchers may be interested in determining the specific amount of data required to reach a desired level of model performance, taking into account the trade-off between costs and outcomes.

To investigate these unresolved questions, we first conducted a systematic survey of 57 published research articles featuring neuroimaging-based predictive models of pain. We also conducted benchmarking analyses on a large-scale functional Magnetic Resonance Imaging (fMRI) pain dataset (*N* = 124), in which we delivered thermal stimuli to induce heat-induced pain and collected pain intensity ratings. Through our literature survey and benchmarking analyses, we explored the impact of various modeling choices on model performance, providing a bird’s eye view of neuroimaging-based pain prediction. This could offer researchers a useful guide for decision-making concerning modeling options, e.g., what could be the modeling targets, which features should be extracted and how they could be combined, and how large the sample size should be, etc. Unlike previous studies that usually examined the impact of each modeling option in isolation, here we systematically compared them all together using a single large-scale pain fMRI dataset. This would offer useful insights for the development of better-performing neuroimaging pain biomarkers and their potential translation into clinical settings, presenting an overarching view of the field of neuroimaging-based pain prediction.

## Results

### Survey results on the use of modeling options

Figure 2 shows the article selection process, and Figures 3–5 show the literature survey results. First, as shown in Figure 3a, the fMRI was the most popular neuroimaging modality for modeling—i.e., 57.9% out of the 57 studies used fMRI. The number of research on neuroimaging-based pain biomarkers has increased since the first publication in 2009. Note that the plot does not display the number of publications of 2020 (*n* = 7) and 2021 (*n* = 1) given that our survey covers only eight months of 2020 (our survey period was between January 2008 and August 2020). Figure 3b shows the study populations of the 57 studies. A greater number of studies addressed clinical pain outcomes (54.4%) as compared to healthy ones (45.6%), and chronic low back pain (cLBP) was the most popular among clinical pain conditions (7 studies). Figure 3c shows the survey results of 157 predictive models from 57 studies. The survey highlights several key trends in the field: (1) Prediction task: Binary classification models (72.6%) were predominant, outnumbering regression models (26.7%) by approximately threefold. (2) Modeling targets: Over half of the models focused on “pain intensity” (27.4%) and “pain vs. no pain” (23.5%). “pain patients vs. controls” was also an important target (21.6%). Note that “pain intensity” included the following two sub-targets, “pain ratings” and “high pain vs. low pain”, which were used for further benchmarking analysis. (3) Model level: A majority of the models were population-level (66.8%), which was almost double the number of idiographic models (31.2%). This result suggests that studies on pain biomarkers usually focus on their clinical applications. (4) Data level: Trial-level (47.7%) and individual-level (33.1%) were most commonly used. (5) Spatial scale: Brain-wide models were more prevalent (56.0%) compared to region-based models, including single-region (26.7%) and multi-region models (17.2%). (6) Experimental task: The phasic pain task was the most common (57.9%), followed by resting-state tasks (24.2%). (7) Feature type: Activation patterns were the most frequently used features (43.9%). (8) Algorithm: Linear SVM was the most utilized algorithm (49.0%), followed by Principal Component Regression with Lasso regularization (LASSO-PCR; 6.3%) and linear SVR (5.7%). Notably, the top three algorithms were all linear methods. (9) Validation methods: Leave-one-out cross-validation was most common (59.8%), with K-fold cross-validation next (17.2%). Figure 3d depicts the distribution of sample sizes across 66 unique datasets. The median sample size was 30, indicative of the typical scale for fMRI experiments.

**Figure 2.**
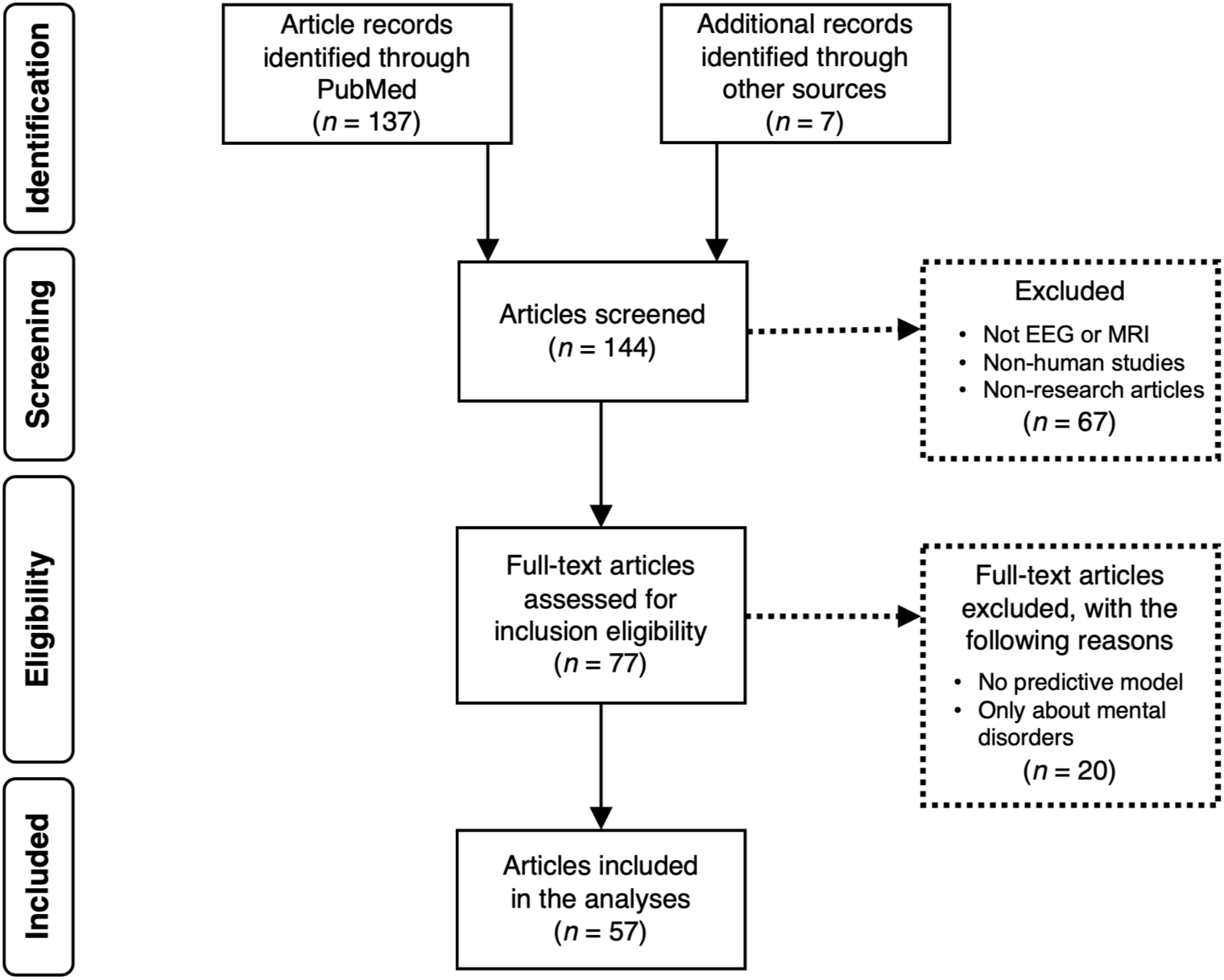
Flowchart of the article selection process for inclusion in the study. This flowchart outlines the systematic approach for selecting articles from initial identification through PubMed and other sources to the final inclusion for model comparison. Research articles from other sources include the articles that were cited by the searched articles but were not identified through the PubMed search. The process includes a comprehensive search, screening of titles and abstracts, eligibility assessment of full-text articles, and the exclusion criteria leading to the final selection. The diagram also shows the numbers of the selected articles for each step. The details of the survey results are shown in **Figs. 3-5**.

**Figure 3.**
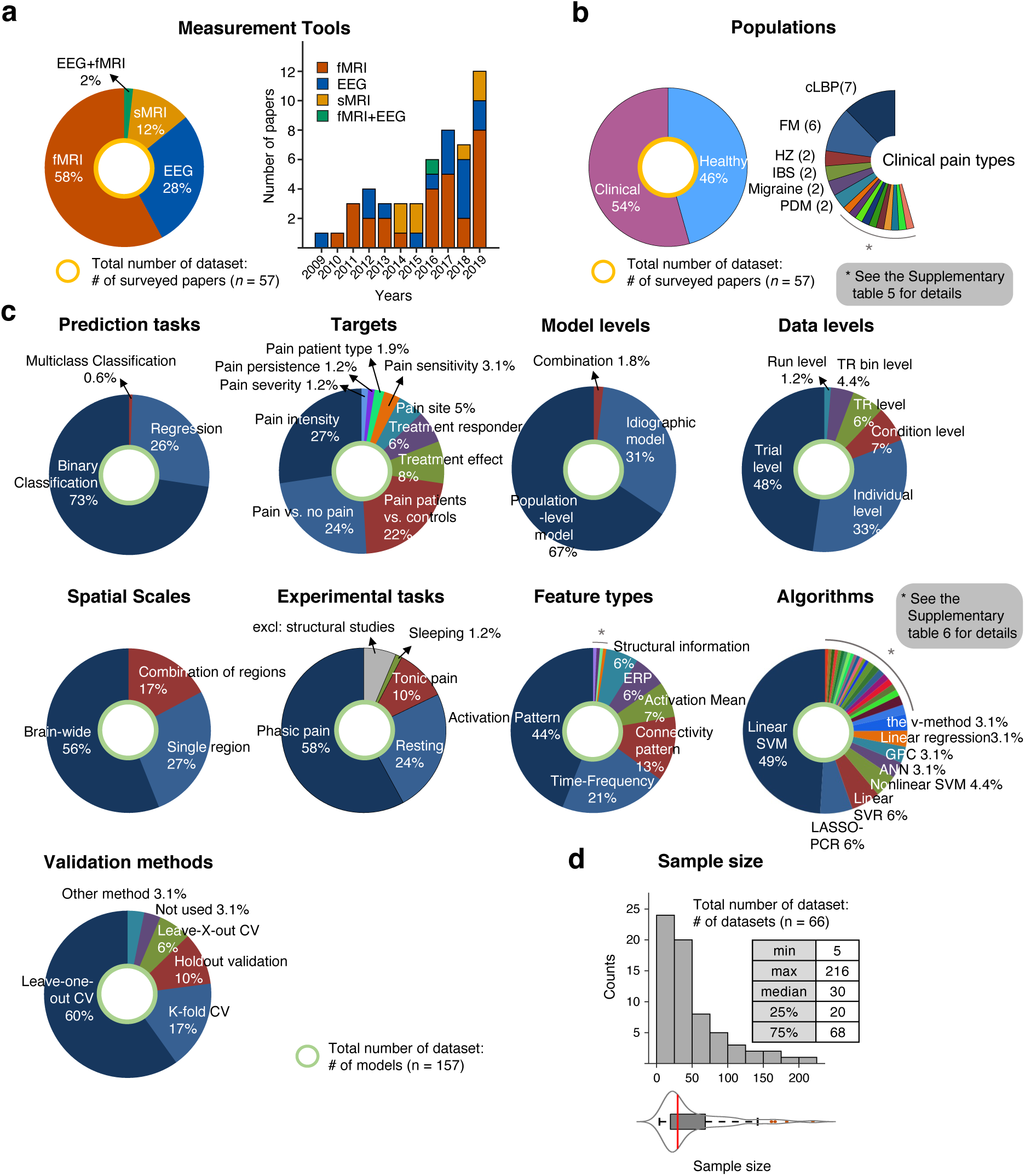
Survey results of neuroimaging-based predictive models of pain. **(a)** This illustrates the distribution of measurement tools used in the final 57 studies selected for the survey, highlighted by the yellow circle. It also shows the distribution of the publication years of these studies. **(b)** The pie charts, based on the same set of studies, detail the proportion of healthy versus clinical populations used to train predictive models and categorize the types of clinical pain investigated. **(c)** Out of the full-text review, 157 models were included for analysis, as indicated by the green circle. The pie charts break down the categories of prediction tasks, model levels, spatial scales, experimental tasks, feature types, validation methods, and algorithms employed. Studies on structural neuroimaging data were omitted from the ‘Experimental tasks.’ **(d)** The box plot represents the sample size distribution across 66 unique datasets from 57 studies. The red line indicates the median value. cLBP, chronic low back pain; FM, fibromyalgia; HZ, herpes zoster; IBS, irritable bowel syndrome; PDM, primary dysmenorrhea; TMD, temporomandibular disorders; SVM, support vector machine; SVR, support vector regression; ANN, artificial neural network; GPC, gaussian processes classification; LASSO-PCR, least absolute shrinkage and selection operator-principal component regression.

### Survey results on model performance

We then compared the model performances across different modeling targets and options. For these comparisons, we focused on the following six aspects—prediction tasks, modeling targets (Figure 4), data and model levels, spatial scale, and sample size (Figure 5). These aspects are among the key aspects that can provide references for future studies and are also included in the later benchmarking analysis.

**Figure 4.**
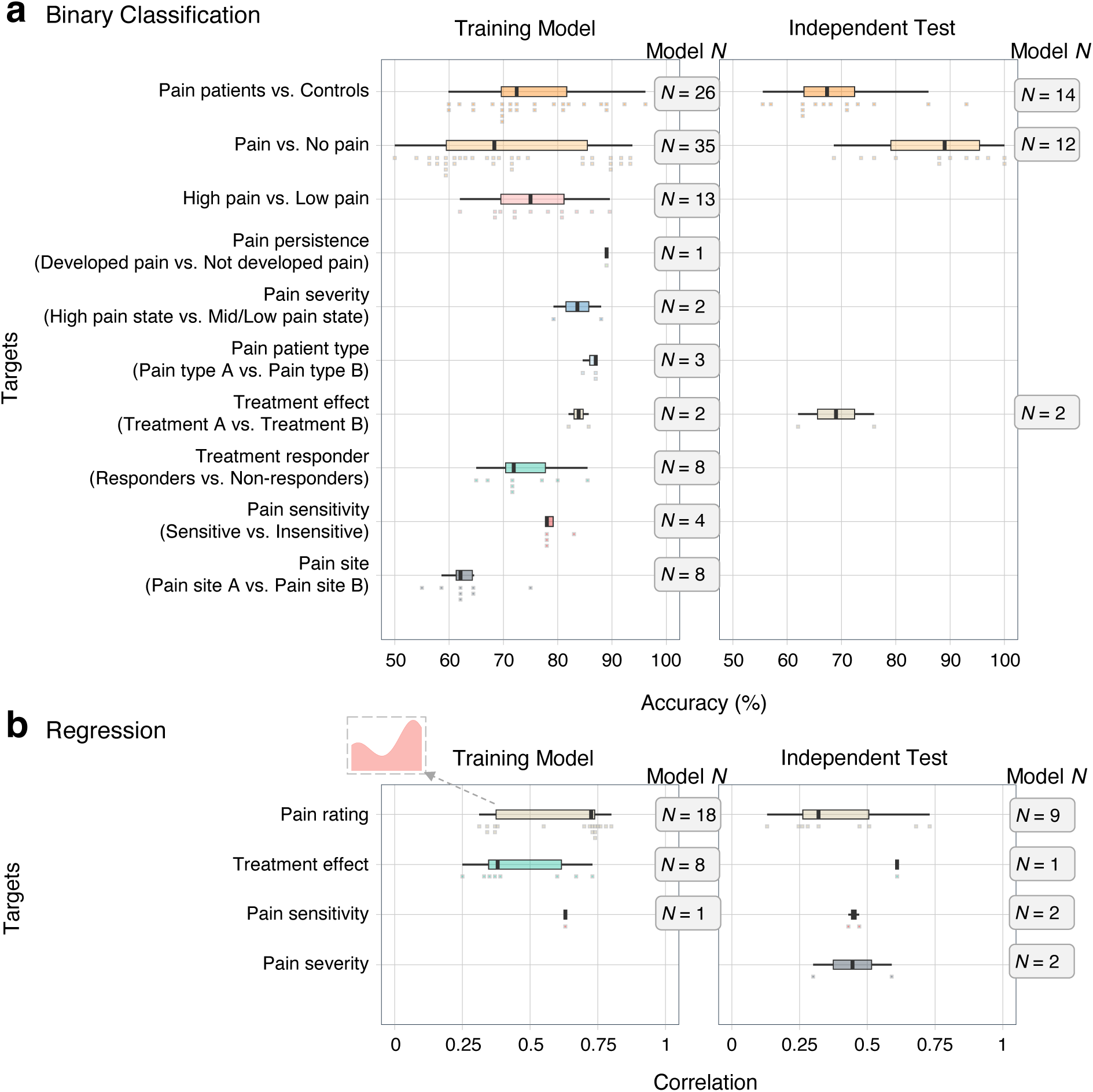
Survey results 1: Model performance across different modeling targets. **(a)** Binary classification task. The box plots illustrate performance distributions for ten different modeling targets during model training and independent testing phases. Each dot represents the classification accuracy (%) of each model. ‘Model *N*’ indicates the number of models included in the comparisons. The targets were ordered by their peak performance values. **(b)** Regression task. Prediction performances of regression-based models were presented for four regression targets. We used prediction-outcome correlation (i.e., a correlation between the predicted and actual values) for comparison. A histogram for ‘Pain rating’ in the training model set is provided, highlighting the bimodal distribution of model performance for the target.

**Figure 5.**
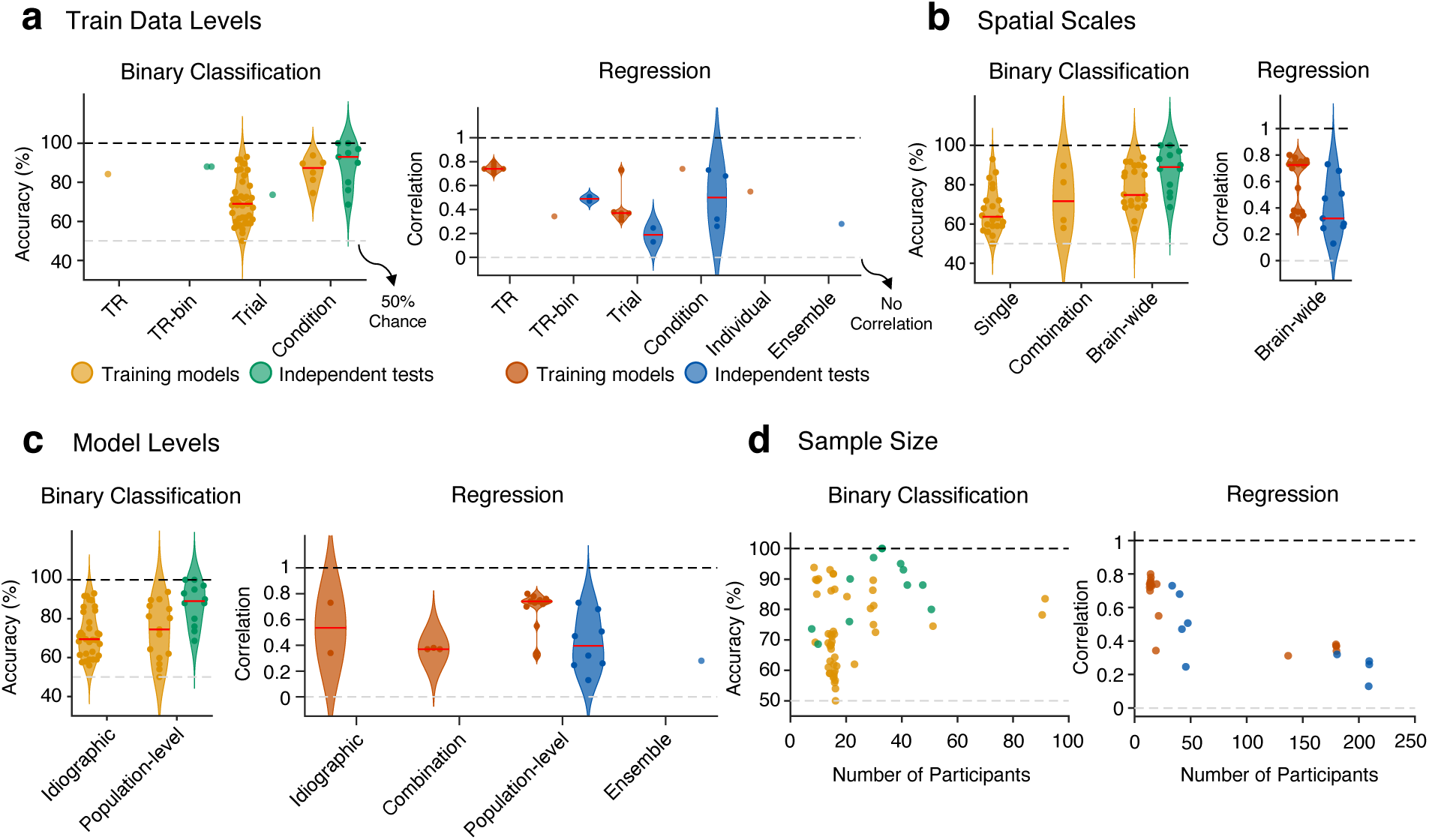
Survey results 2: Model performance across different data levels, spatial scales, model levels, and sample sizes. **(a-c)** Violin plots illustrate the distribution of model performance across various aspects: training data levels **(a)**, spatial scales **(b)**, and model levels **(c)**. In these plots, individual dots represent the performance of each model, quantified as accuracy for binary classification tasks or prediction-outcome correlation for regression tasks. Yellow and red plots represent the performance derived from the training datasets, while green and blue plots illustrate the performance observed in independent tests. The median performance for each category is indicated by red lines within the plot. A black dashed line represents the possible maximum performance, and a gray dashed line denotes the baseline level of performance, which corresponds to chance or no correlation. **(d)** Impact of sample size on model performance. Scatter plots demonstrate how the number of participants in a model influences performance, with separate visualizations for binary classification and regression tasks. Yellow and red dots represent the performance derived from the training datasets, while green and blue dots denote the performance in independent tests.

First, we compared the classification accuracy and prediction-outcome correlation (the correlation between predicted and actual values) across various modeling targets. To present these results effectively, the plots in Figure 4 are organized by the maximum performance values achieved by the training models for each specific target. The most frequently studied targets were “Pain patients vs. Controls” (26 training models & 14 independent tests), “Pain vs. No pain” (35 training models & 12 independent tests), and “Pain rating prediction” (18 training models & 9 independent tests). “High pain vs. Low pain” was also a popular target, but there was no independent test (13 training models & no independent test). “Treatment responder,” “Treatment effect,” and “Pain site” were the next popular targets, but these also had one or no independent test, highlighting that these targets need further validation studies. For “Pain patients vs. Controls” and “Pain rating prediction,” the model performances for the independent tests (median accuracy = 67.3%, median *r* = 0.32) were lower than the training results (accuracy = 72.4%, *r* = 0.725), indicating the potential bias in the reported model performances. For “Pain vs. No pain,” some independent tests showed higher performances than training, which was from one specific study that showed high accuracy in multiple independent tests (Wager et al., 2013). Note that targets with a large number of reports showed high variance in their model performance, suggesting that the targets with a small number of reports may not be able to serve as references for future studies and require further studies. In addition, the model performance of “Pain rating prediction” exhibited a bimodal distribution in training results (Figure 4b). This, along with the high variance in the classification model performances, implies that other variables and modeling options may have significant impacts on model performance.

We then evaluated the influences of modeling options on the model performance, as shown in Figure 5. Here are some observations. First, the survey on the train data levels showed that the condition level results were higher than the trial level results (median accuracy = 87.3% for the condition level training models, and 69.0% for the trial level training models; Figure 5a). Second, also from Figure 5a, while classification models did not exhibit significant decreases in performance in independent tests, regression models showed an overall decline in performance, suggesting that regression models may have more difficulty in generalizing than classification models. This trend was also observed in the relationship between model performance and sample sizes in Figure 5d. Third, for the spatial scale, as shown in Figure 5b, the brain-wide models showed higher performance (median accuracy = 74.8% for brain-wide training models) than a combination of regions (71.6%) or single region (63.7%). Lastly, for the model levels (Figure 5c), the population-level models showed the highest performance both in classification and regression models (median accuracy = 74.5% and median *r* = 0.74 for population-level training models).

### Benchmarking analysis

Given that the different experimental designs and populations across surveyed studies limited the direct comparability of the results, we additionally performed benchmarking analyses on the locally collected large-scale pain fMRI dataset (*n* = 124) with a single experimental design. Our benchmarking analysis focused on the following four aspects: (1) Data level, (2) Spatial Scale, (3) Model Level, and (4) Sample Size, and two different targets: binary classification of high versus low pain and the prediction of pain ratings with regression models. Figure 6 summarizes the benchmarking analysis pipeline. Importantly, all the results presented here were obtained from tests on the hold-out test set of 44 participants.

**Figure 6.**
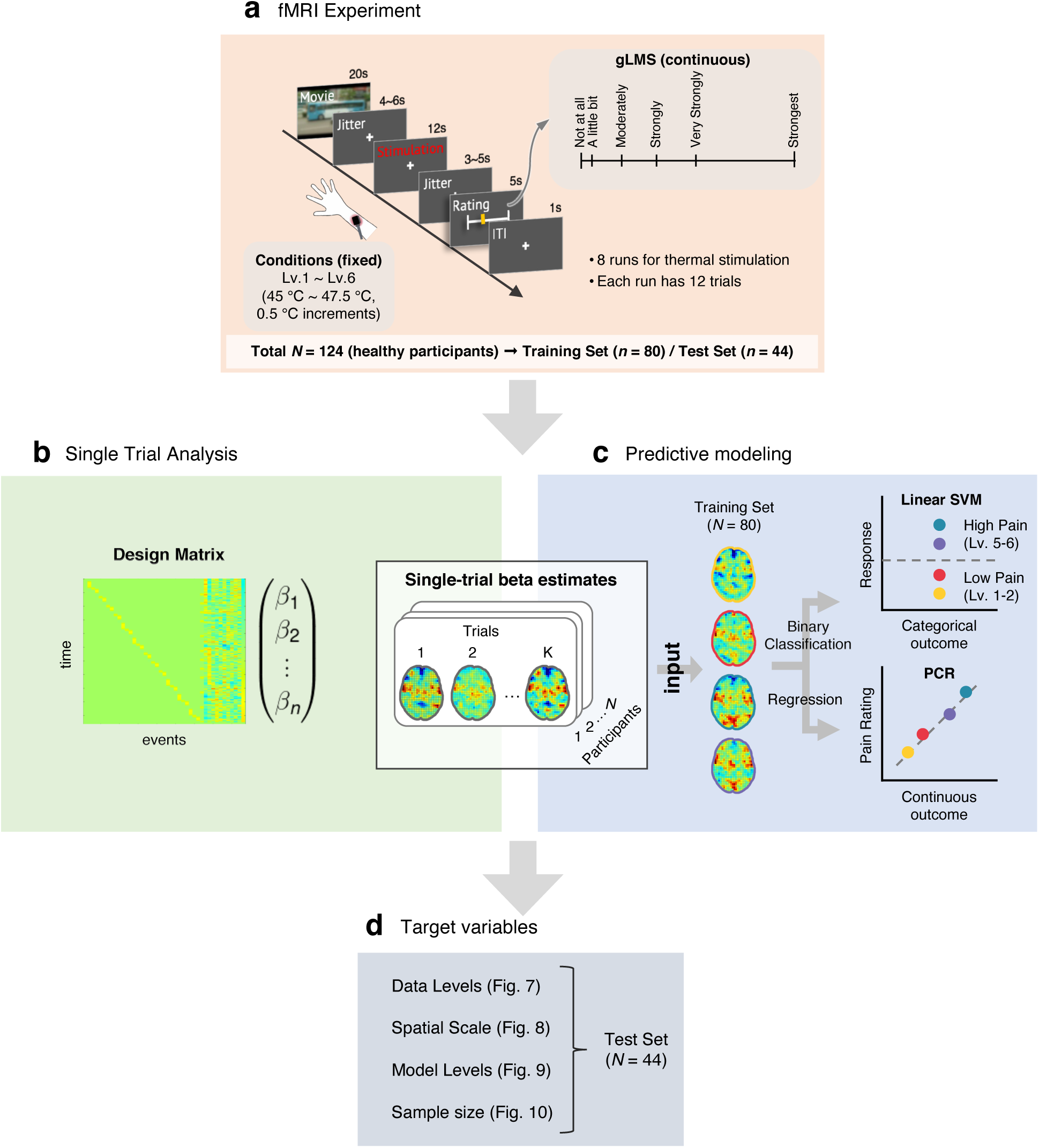
Overview of Benchmarking Analysis. We performed a benchmarking analysis to examine the impact of modeling options on model performance. **(a)** fMRI experiment setup. We collected task-based fMRI data from 124 healthy participants. We delivered thermal stimulation with six levels of stimulus intensity, ranging from 45 to 47.5 ℃ in 0.5 ℃ increments. After the heat stimulation, participants were asked to rate pain intensity on the generalized Labeled Magnitude Scale (gLMS) (Bartoshuk et al., 2004). Participants completed a total of eight runs of thermal stimulation, two of which were without a movie and six runs with a movie. In the case of runs with a movie, a 20-second movie clip was shown before the thermal stimulation. Each run consisted of 12 trials. For model training and testing, we divided the dataset into a training set (*N* = 80) and an independent test set (*N* = 44). All results were based on the independent test dataset. **(b)** Single-trial analysis. We obtained single-trial voxel-wise beta maps for each participant using a general linear model (GLM) with separate regressors for each trial, as in the “beta series” approach (Rissman et al., 2004). These beta maps served as inputs for subsequent analyses. **(c)** Predictive modeling. We develop predictive models with machine learning techniques. For binary classification, the target was ‘high’ versus ‘low’ pain. We defined “high” pain as heat stimulus levels 5 and 6 and “low” pain as heat stimulus levels 1 and 2. For binary classification, we used linear Support Vector Machines (Linear SVMs). For regression-based prediction, the target was ‘pain ratings,’ and we used Principal Component Regression (PCR) for model training. **(d)** Target variables. We provide results about four aspects (i.e., data levels, spatial scales, model levels, and sample size). The details of the results are shown in the following figures (**Figures 7–10**).

#### Benchmarking analysis (1): Data level

The first benchmarking analysis was on the data level. The “data level” indicates the number of trials averaged for model training and testing (Figures 7a-b). Figure 7c provides the classification model performance (i.e., accuracy), and Figure 7d provides the regression model performance (i.e., prediction-outcome correlation). In Figures 7c-d, the plots in the first and second columns present the benchmarking analysis results for the train vs. test data levels, respectively, focusing on the impact of different train vs. test data levels on model performance. The plots in the third column show the benchmarking analysis results for the test data levels but with the matched number of test data, which was 6 data points.

**Figure 7.**
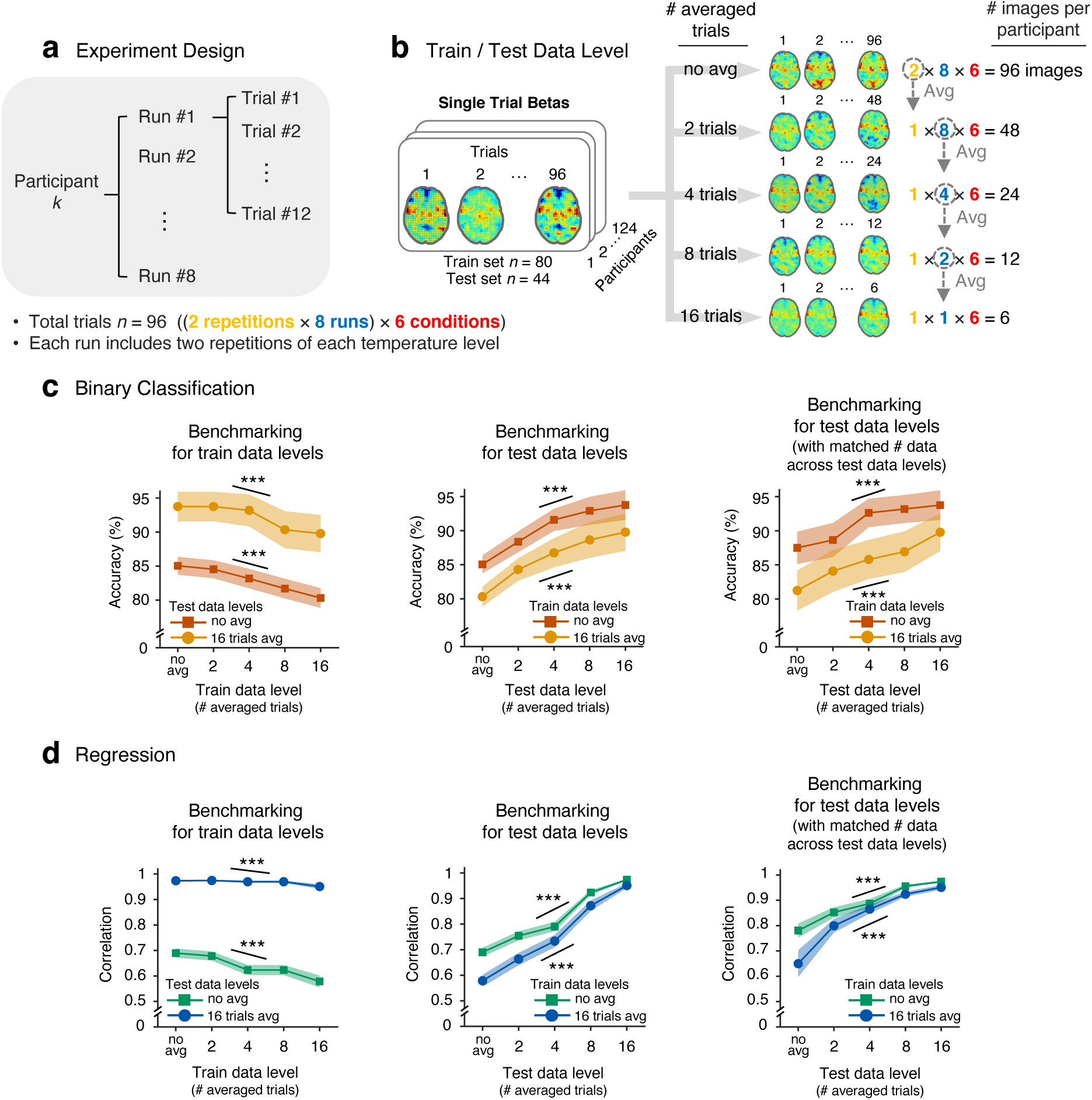
Benchmarking analysis results for data levels (i.e., data averaging). **(a)** Data structure. Each participant’s data consisted of 96 trials, distributed over 8 runs, with each run comprising 12 trials across 6 temperature conditions, with each temperature level repeated twice. **(b)** Data were analyzed at five averaging levels: single trials (no averaging), and averages of 2, 4, 8, and 16 trials. This averaging was applied separately to both training and testing datasets. **(c-d)** The plots show the impact of averaging on model performance in the binary classification (c) or the regression (d) tasks. **Left**: The effect of training data averaging on model performance at two test data levels (‘no averaging’ and ‘16 trials averaged’), **Center**: The effect of test data averaging on model performance at two train data levels (‘no averaging’ and ‘16 trials averaged’), **Right**: The effect of test data averaging on model performance at two training data levels, with a fixed number of images in the test set (6 data points). The lines indicate mean accuracy or correlation (respectively) across 44 participants, with shadings representing the standard error of the mean. The color scheme differentiates the averaging conditions with red/green indicating no averaging and yellow/blue indicating 16 trials averaged; we selected these two data levels as they were the most commonly used based on our literature survey (i.e., single-trial and condition level). Model performance exhibits significant trends with respect to data averaging, as established by a multi-level GLM analysis. Detailed performance metrics and statistical results are provided in **Supplementary Table 1**. \*\*\**p* < 0.001.

For the high versus low pain classification, we found that model performance decreased as more data were averaged in the training dataset (Figure 7c). For example, when the test data level was ‘no average,’ the model performances across 5 different train data levels were 85.05% ± 1.31% (mean accuracy ± s.e.m.) for ‘no average’, 84.53% ± 1.32% for two trials averaged, 83.17% ± 1.43% for four trials averaged, 81.70% ± 1.44% for eight trials averaged, and 80.31% ± 1.45% for 16 trials averaged. The same patterns were also observed in the other test data level (i.e., 16 trials averaged). A multilevel general linear model (GLM) showed that the trend of decreasing model performance over averaging the training data was significant, beta = −1.09, *z* = −3.53, *p* = 0.00042, two-tailed, bootstrap test. The same pattern was also observed when 16 trials were averaged for the testing data, beta = −5.09, z = −4.50, *p* = 0.00001, multi-level GLM, two-tailed, bootstrap test.

In contrast, we observed that the model performances increased as more data were averaged in the testing dataset. For example, when the train data level was fixed at ‘no average’, the model performances across 5 different test data levels were 85.05% ± 1.31% (mean accuracy ± s.e.m.) for ‘no average’, 88.36% ± 1.52% for two trials averaged, 91.57% ± 1.62% for four trials averaged, 92.90% ± 2.05% for eight trials averaged, and 93.75% ± 2.17% for 16 trials averaged. A multilevel GLM showed that the trend of increasing model performance over averaging the testing data was significant, beta = 2.35, *z* = 3.43, *p* = 0.00061, two-tailed, bootstrap test. The same pattern was observed for 16 trials averaged for the training data, beta = 2.28, *z* = 3.40, *p* = 0.00068, multi-level GLM, two-tailed, bootstrap test. Furthermore, the same trend was observed when we compared the performance across 5 different test data levels with a matched number of test data points (i.e., 6 data points per participant) for both ‘no average’ and 16 trials averaged for the training data, betas = 4.56 and 3.49, *zs* = 3.78 and 3.18, *ps* = 0.00015 and 0.00147, multi-level GLM, two-tailed, bootstrap test.

Lastly, the main findings for the pain rating regression were similar to the binary classification results—the model prediction performance increased as more data were averaged in the test dataset, while it decreased as more data were averaged in the training dataset. These effects were significant and consistent across different combinations of train and test data levels (Figure 7d). For example, multi-level GLMs with a linear regressor revealed significant increases in model performance across different test data levels for all train data levels—‘no average’ (beta = 0.07, *z* = 4.43, *p* = 0.00001, two-tailed, bootstrap) and ‘16 trials averaged’ (beta = 0.09, *z* = 3.93, *p* = 0.00009). The details of the model performance and significant test results are in **Supplementary Table 1**.

#### Benchmarking analysis (2): Spatial scale

The second benchmarking analysis was on the spatial scale. Figure 8a presents an overview of the analysis. While increasing spatial scales, we developed new models and calculated the classification accuracy (Figure 8b) and prediction-outcome correlation (Figure 8c) based on the test dataset (*n* = 44). The model development was repeated 100 times using random combinations of brain regions. For “brain-wide masks,” we employed three distinct masks: one encompassing 21 pain-predictive regions (‘21’) from a previous study (Kohoutova et al., 2022), the Neurosynth ‘Pain’ mask (‘NP’), and a gray matter mask (‘GM’) without iteration. Figures 8b-c show model performance across 44 participants in the independent dataset.

**Figure 8.**
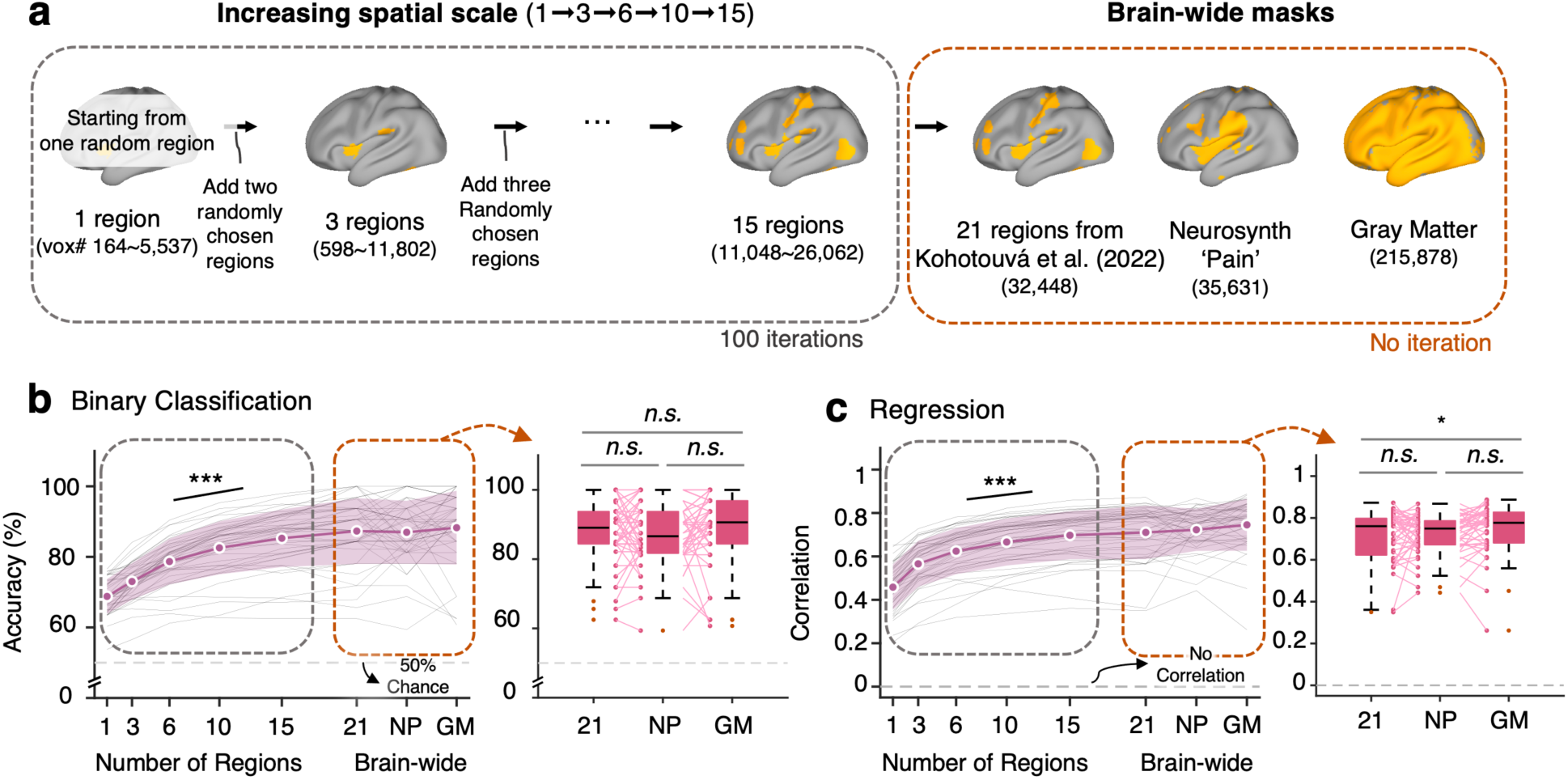
Benchmarking analysis results for spatial scales. **(a)** The analysis employed two different approaches. In the first approach, we used a progressive mask construction method. Starting with one randomly selected region, we incrementally added more regions, up to a total of 15. These regions were selected from predefined regions-of-interest (ROIs) identified in a previous study (Kohoutova et al., 2022). Predictive models were then trained using these incrementally constructed masks. This iterative process was repeated 100 times, as indicated in the illustration by a gray dashed line box. The second approach involved the use of three comprehensive brain-wide masks. These included: (1) a mask encompassing all 21 ROIs from (Kohoutova et al., 2022), (2) a ‘Pain’ mask derived from Neurosynth (labeled as NP), and (3) a gray matter mask (GM). Each mask’s voxel count is noted below its respective label. For the cumulative regions mask, a range is provided instead of a fixed number, reflecting the variability due to the random selection process. This approach is delineated in the box with an orange dashed line. **(b-c)** The average performance from 100 iterations was calculated for each spatial scale. The reddish-purple lines and shaded areas depict the mean and standard deviation of performance, respectively, across 44 participants in the test dataset. Gray thin lines show model performance for each participant. Performance trends showed a significant increase with the number of regions, validated by multi-level general linear models (GLMs) with *z*s = 3.66 and 3.58, *p*s = 0.00025 and 0.00034 for binary classification (b) and regression (c), respectively, two-tailed, bootstrap tests. The boxplots show the differences in model performance among the three brain-wide masks. *n.s. p* > 0.05, \**p* < 0.05, \*\*\**p* < 0.001. NP, Neurosynth pain mask; GM, Gray matter.

The results indicated that the model performance increased with the increasing numbers of combined regions. For the increasing numbers of combined regions (i.e., 1, 3, 6, 10, and 15 regions), the mean binary classification accuracies were 68.79%, 73.01%, 78.67%, 82.56%, and 85.27% (Figure 8b and **Supplementary Table 2**, and the prediction-outcome correlations were 0.46, 0.57, 0.62, 0.67, and 0.70 (Figure 8c and **Supplementary Table 2**). These increasing trends of model performance were statistically significant, betas = 4.25 and 0.06, *z*s = 3.66 and 3.58, *p*s = 0.00025 and 0.00034, two-tailed, bootstrap test, multi-level GLM. For the tests for “brain-wide masks,” there were no significant differences among the type of brain masks for the binary classification task, 87.31%, 86.96%, and 88.27% for ‘21’, NP, and GM, respectively, all |*t|*s < 0.98, *p*s > 0.33, two-tailed, paired *t*-test. However, there was a significant difference between GM versus ‘21’ for the rating prediction task, prediction-outcome *r* = 0.75 (GM) and 0.71 (‘21’), *t*(43) = 2.626, *p* = 0.0119, two-tailed, paired *t*-test, suggesting that the information distributed across the gray matter, even outside of the 21 pain-predictive regions, was helpful for predicting pain ratings. Lastly, there were no significant differences between ‘21’ versus NP and GM versus NP for the rating prediction task, prediction-outcome *r* for NP = 0.72, *t*(43) = 0.764, *p* = 0.4489 for ‘21’ versus NP, *t*(43) = 1.962, *p* = 0.0562 for GM versus NP.

#### Benchmarking analysis (3): Model level

The third benchmarking analysis was on the model level. The “model level” analyses compared the test results between the idiographic versus population-level predictive modeling (Figure 9a, for details, see **Materials and Methods**). Briefly, idiographic predictive modeling indicates the modeling based on a single participant’s data, whereas population-level predictive modeling indicates the modeling using the data combined across all individuals. There were also two types of independent tests in this benchmarking analysis. First, we tested the models on the within-individual hold-out data of the training dataset (*n* = 61). For this, we held out two-run data for the testing by using only six-run data for training. Second, we tested the models on the fully independent test dataset (*n* = 44). We combined the different training and testing methods, which resulted in the following four combinations: ① idiographic model (‘Id’) tested on within-individual hold-out data, ② population-level model (‘Po’) tested on within-individual hold-out data, ③ averaged idiographic model (‘Avg Id’) tested on the independent test dataset, and ④ population-level model (‘Po’) tested on the independent test dataset. Figures 9b-c show the comparisons of the classification and the regression model performances. We conducted two-sample *t*-tests to compare model performance between the idiographic versus population-level models.

**Figure 9.**
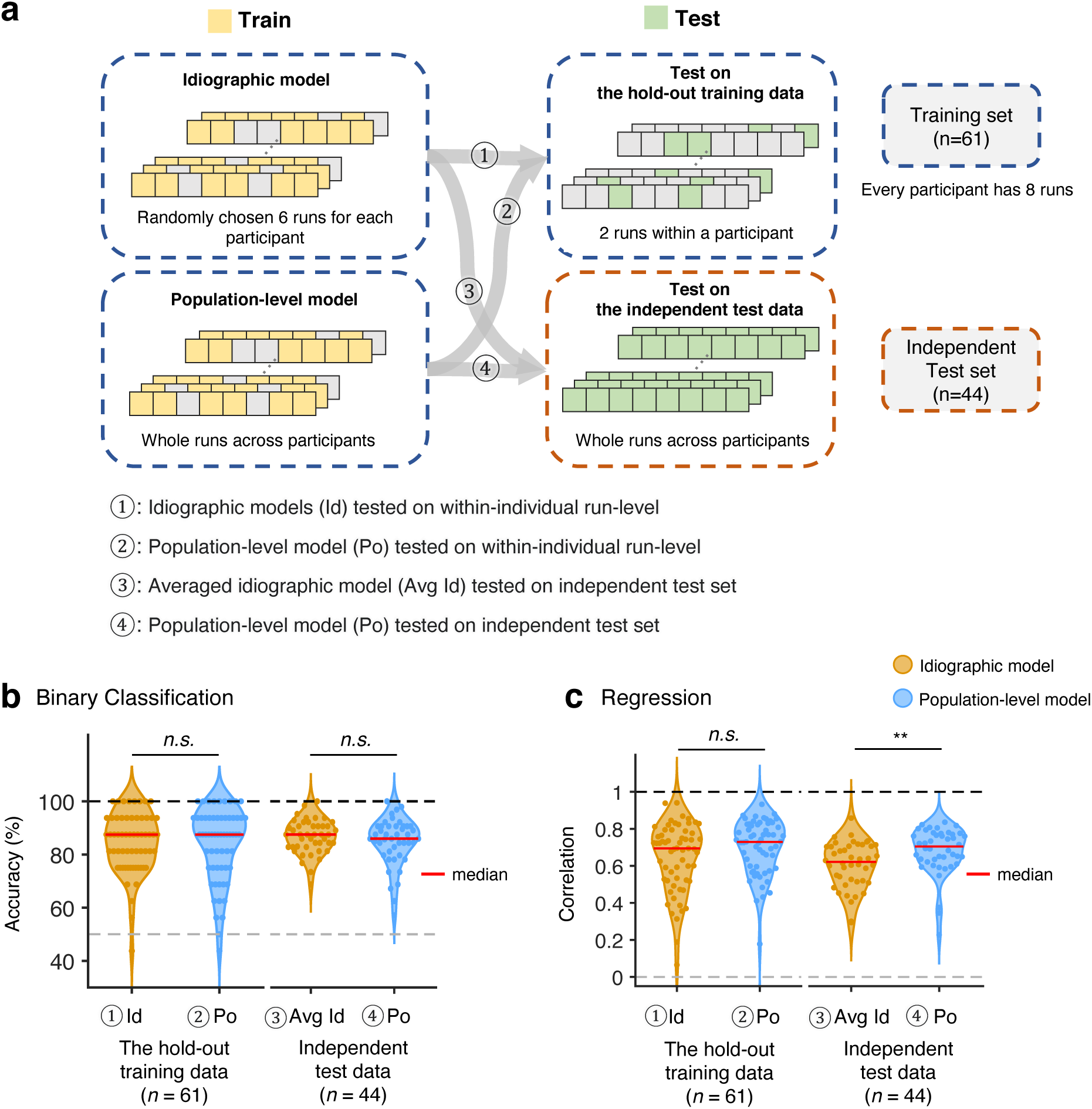
Benchmarking analysis results for model levels. We provide a benchmark analysis of model performance at the individual (idiographic) and group (population) levels. In this analysis, we used data from 6 runs for training and data from 2 runs for testing. This choice required the use of data from only 61 participants who had complete data for all 8 runs. **(a)** Model training and testing procedures. Idiographic model (Id): Predictive models were trained on trial-level data from six runs per participant and tested on unseen data from the remaining two runs (①). In addition, we created a group-level model by averaging all individual models and tested the model on an independent test dataset (③). Population-level model (Po): Models were developed using trial-level data from all participants in the training dataset. Again, the data from six runs were used. The model was tested on both unseen data from the remaining two runs of 61 participants (②) and data from an independent test dataset (④). **(b)** Binary classification. No significant performance differences were found between the idiographic and population-level models when evaluated on both the hold-out training data and the independent test dataset (*p*-values = 0.7868 for ① vs. ② and 0.2358 for ③ vs. ④, two-sample *t*-test). **(c)** Regression. There was no significant difference in performance on the hold-out training data (*p*=0.2005 for ① vs. ②); however, the population-level model significantly outperformed the averaged idiographic model on the independent test dataset, *p*=0.0061 for ③ vs. ④, two-sampled *t*-test. \*\**p* < 0.01.

In the binary classification, the performance of idiographic and population-level models did not significantly differ, *t*(120) = 0.271, *p* = 0.7868 for testing on the within-individual hold-out data, *t*(86) = 1.194, *p* = 0.2358 for testing on the independent test dataset, two-tailed, two-sample *t*-test. In the regression analysis, while the idiographic and population-level models demonstrated comparable performance on within-individual hold-out data, *t*(120) = 1.287, *p* = 0.2005, the population-level model significantly outperformed the idiographic model on the independent test dataset, *t*(86) = 2.813, *p* = 0.0061, two-tailed, two-sample *t*-test. **Supplementary Table 3** provides detailed results of model performance.

#### Benchmarking analysis (4): Sample size

The final benchmarking analysis focused on the impact of sample size (**Figure 10a**). While increasing the sample size, we developed new models and calculated the classification accuracy (**Figure 10b**) and prediction-outcome correlation (**Figure 10c**) based on the independent test dataset (*n* = 44). The model development was repeated 100 times using a random selection of participants. (“Increasing sample size” in **Figure 10a**). For “full sample size,” all 80 participants from the training dataset were used without the random selection (and thus no iteration). The model performances on the independent test dataset (*n* = 44) are shown in Figures 10b-c.

**Figure 10.**
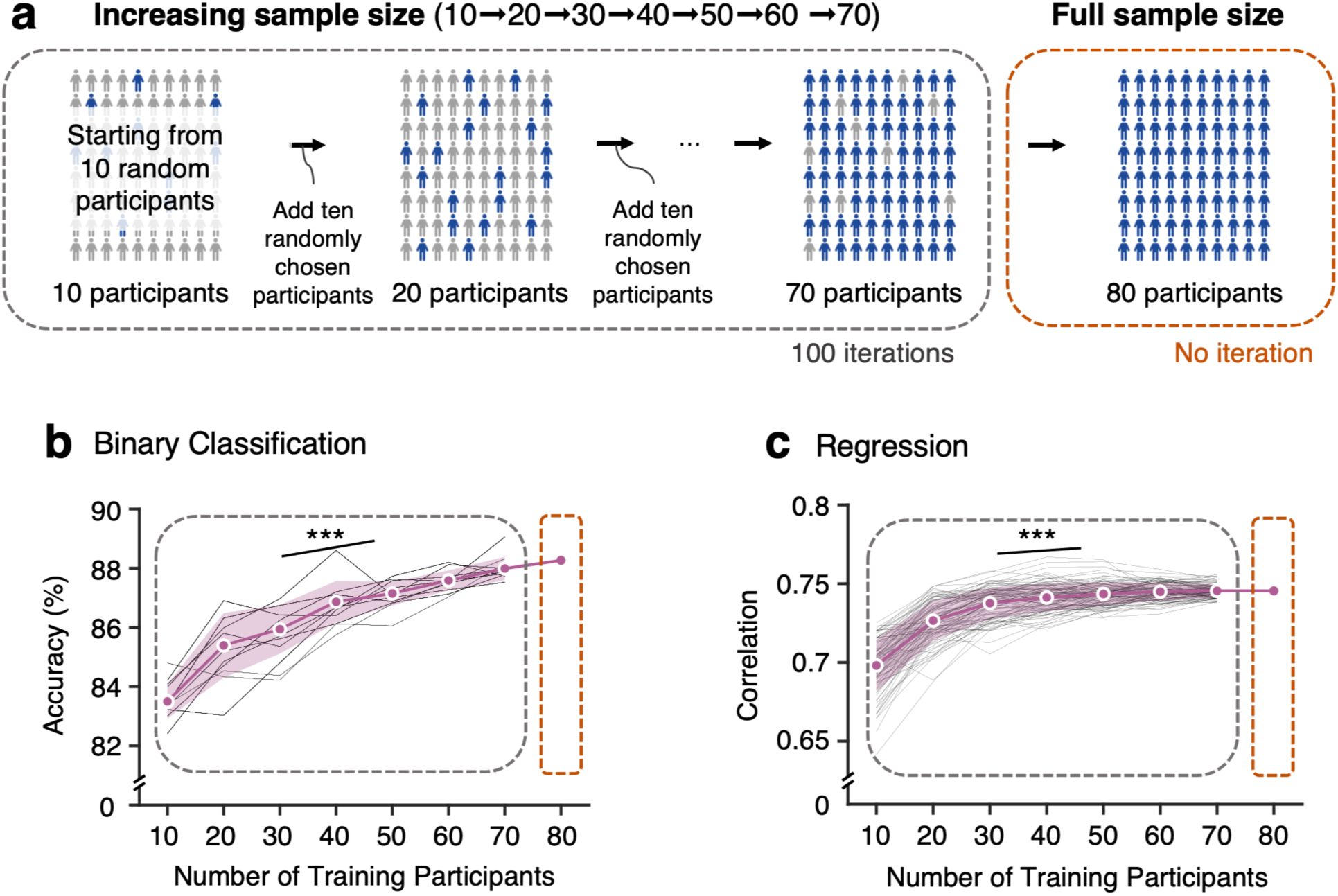
Benchmarking analysis results for sample size. We investigated how the number of participants in the training set affects the performance of predictive models. Models were iteratively developed with increasing training sample sizes, ranging from 10 to 70 participants, and compared to a model trained with the full set of 80 participants. **(a)** The study began with 10 randomly selected participants from the training dataset and incrementally and randomly added sets of 10 participants to build new models. This incremental process was repeated 100 times, as indicated by the box with a gray dashed line. In a separate analysis, the full cohort of 80 participants was used, as indicated by the box with an orange dashed line. **(b-c)** Average model performance across iterations is shown by the reddish-purple line, with the shaded area indicating the standard deviation, for the binary classification (b) and regression (c) tasks. Gray thin lines show model performance in each iteration. Statistically significant improvements in model performance were observed as the number of training participants increased (for the binary classification and regression tasks, *z*s = 3.96 and 4.02, *p*s = 0.00008 and 0.00006, respectively, two-tailed, bootstrap test. The full sample size models performed better than the other models with smaller sample sizes, achieving 88.26% accuracy for binary classification and a correlation of 0.74 for regression. \*\*\**p* < 0.001.

The test results for the “increasing sample size” condition showed that the model performance increased with the increasing numbers of training samples. The binary classification accuracies (mean ± standard deviation) for the increasing numbers of combined samples (i.e., 10, 20, …, 60, and 70 participants) were 83.50% ± 0.58, 85.39% ± 1.09, 85.94% ± 0.83, 86.86% ± 0.70, 87.15% ± 0.40, 87.59% ± 0.34, 87.99% ± 0.40 (**Figure 10b** and **Supplementary Table 4**), and the prediction-outcome correlations were 0.698 ± 0.01, 0.727 ± 0.01, 0.738 ± 0.01, 0.741 ± 0.01, 0.744 ± 0.01, 0.745 ± 0.01, and 0.746 ± 0.004 (**Figure 10c** and **Supplementary Table 4**). These increasing trends of model performance were statistically significant (betas = 0.68 and 0,01, *z*s = 3.96 and 4.02, *p*s = 0.00008 and 0.00006, two-tailed, bootstrap test, multi-level GLM).

## Discussion

In this study, we conducted a literature survey and four benchmarking analyses on neuroimaging-based pain biomarkers. The importance of building pain biomarkers lies in the need for objective assessment tools for pain because of the subjective nature of pain assessment, which hampers the choice of appropriate interventions or the development of new treatments for pain. A recent study conducted a qualitative review on the topic (van der Miesen et al., 2019), but more systematic analyses on the choices of modeling targets and options have yet to be done. To fill the gap, we systematically compared the performance of different models from previously published studies on neuroimaging-based pain biomarkers and conducted benchmarking analyses using a large-scale fMRI pain dataset (*N* = 124). In these analyses, we focused on the following modeling aspects—prediction tasks, modeling targets, data levels, spatial scales, model levels, and sample sizes.

Our survey results provide multiple interesting observations. First, the survey revealed that there were a few preferred targets and options for predictive modeling. The majority of the predictive models were designed for classification (73%), aimed at population-level prediction (66%), used brain-wide features (56%), and were trained on the data at the trial- or individual-level (80%). These characteristics suggest that those who developed the models were mindful of the models’ clinical applications and their utility (Davis et al., 2020) because population-level diagnostic models that can be applied to new individuals are favorable for clinical purposes. Second, fair comparisons of model performance across different modeling targets were quite challenging due to the small number of studies for each target. Even when there were many studies for the modeling target, it showed highly variable model performance, suggesting that there were many factors that influenced the level of model performance. Thus, we found that it was not simple to determine the level of difficulty for different modeling targets. To address this issue, we would need more research and systematic investigation of the effects of modeling targets and options. In addition, consideration should be given to the biomarker categories based on their intended use. The U.S. Food and Drug Administration identified seven major categories of biomarkers, which include safety, response, monitoring, diagnostic, predictive, risk, and prognostic biomarkers (Kragel et al., 2021; Mackey et al., 2019; Reckziegel et al., 2019; Tracey et al., 2019). It is likely that the levels of difficulty are varied across these biomarker categories, and thus, the standards for clinical translation should be carefully determined based on future studies. The third observation from the survey pertains to the limited number of independent tests. Only a small proportion of studies (28%) assessed their models on independent datasets, making it challenging to assess the impacts of modeling targets and options on model performances in an unbiased way.

Our benchmarking analysis results also offer several interesting observations regarding the effects of modeling options on model performance. First, the distinct effects of data averaging on model performance were observed for training versus testing data—less data averaging was helpful for training data, whereas more data-averaging was helpful for testing data. This can be understood as a tradeoff between increasing the signal-to-noise ratio by averaging versus expanding the sample distribution. That is, expanding the sample distribution of training data improves model performance, and reducing the noise of testing data also improves model performance. Second, both survey and benchmarking analysis results failed to show the benefits of idiographic modeling compared to population-level models. This is somewhat contradictory to the common notion that personalized models should perform better than population models because the idiographic approach removes one important source of variance—between-individual variability (Lindquist et al., 2017). However, when we used a subset of data from a single session for model training, the idiographic models did not perform better than the population-level models. Considering the recent suggestions about the extensive sampling of small *N* participants (Gratton et al., 2022; Naselaris et al., 2021), the non-significant results may be due to the small amount of data per individual in our sample. Thus, future studies should examine the effects of idiographic versus population-level modeling with multi-session data from a small number of participants. Third, our results showed that including more brain regions and increasing sample sizes improve model performance, consistent with previous studies (Brodersen et al., 2012; Kragel et al., 2018; Marek et al., 2022). However, in the benchmarking analysis, the results suggested that including extra brain regions beyond the regions known to be important for pain prediction did not always guarantee higher model performance. For example, though the number of voxels within the gray matter mask (*n*_vox_ = 215,878) was almost seven times greater than 21 *a priori* pain-predictive regions (*n*_vox_ = 32,448) or Neurosynth mask (*n*_vox_ = 35,631), the classification performance of the gray matter mask-based model was not significantly better than the models based on the 21 pain-predictive regions and Neurosynth mask. However, when it comes to the regression task, the model based on the gray matter mask showed significantly better performance than the 21 region-based regression model, suggesting that the characteristics of the prediction task can affect the benefit of including more regions and voxels. Considering that the test dataset size could also be a factor influencing the task difficulty, future studies should examine the impact of the spatial scale with larger sample sizes in the test dataset.

There are multiple limitations in this study. First, our survey results were limited by the small number of pain biomarker studies. For example, only 28% of the surveyed studies reported test results with independent test datasets, which made it difficult to obtain unbiased estimates of model performances. This means that our survey results could be biased. In addition, due to the insufficient number of studies in the survey, we had to combine results from healthy and clinical populations, though their brain conditions are substantially different (Apkarian et al., 2005). Second, some studies with more than one predictive model could have a greater influence on the survey results. When we compared model performance, we compiled the testing results at the model level but noted that some studies provided multiple models while others provided only a single model. Thus, to take this data structure into account, a hierarchical approach to performance comparisons should be considered in future studies. **Supplementary Table 5** provides the number of models and independent tests for each study. Third, we had to use the most popular performance measures, such as correlation, to compare model performances, but it is well-known that correlation as a performance measure has some caveats (e.g., being insensitive to scaling, providing biased results, etc.) (Poldrack et al., 2020). As more researchers adopt better practices in predictive modeling, we should be able to use better performance measures, such as *R^2^*, for the comparisons. Fourth, in the benchmarking analyses, we did not cover all the aspects considered in the survey due to the limitations of our dataset. For example, our dataset only included fMRI data from healthy participants, and thus, we could not provide results on other neuroimaging modalities or clinical samples. Thus, one should be cautious about generalizing our results to other modalities or clinical samples.

In conclusion, this study investigated the influences of modeling options and targets on the performance of neuroimaging-based pain biomarkers. Through a literature survey and benchmarking analyses, we found that data levels, spatial scales, and sample sizes were important determinants of classification and prediction performance. To improve model performance, incorporating a larger number of pain-related brain regions, increasing the sample sizes, and reducing data averaging in the training dataset while increasing it in the test dataset appeared to be helpful. These findings will serve as a useful reference for making decisions on neuroimaging-based biomarker development, highlighting the importance of a careful selection of modeling variables to build better-performing neuroimaging pain biomarkers.

## Materials and Methods

### Literature survey

We conducted a literature survey to examine the current state of neuroimaging-based pain biomarker research. For more focused analyses, we only included the papers that used MRI or electroencephalogram (EEG) for pain prediction. Figure 2 shows a flow diagram of the article search and inclusion. We conducted a search using PubMed for research articles published between January 2008 and August 2020 with the following search terms: “pain” in the Title/Abstract; “predict” or “classf*” in Title/Abstract; “eeg”, “fmri”, “magnetic resonance imaging”, or “brain” in Title/Abstract; “machine learning”, “predictive modeling”, “decoding”, “signature”, “svm”, or “biomarker” in Title/Abstract; NOT “review” in Publication Type; NOT “Symposium” in Title/Abstract. The number of initially searched articles was 137. The exclusion criteria included (a) non-human animal studies, (b) absence of prediction models, (c) studies not about pain, (d) non-empirical research articles (e.g., review articles), and (e) studies using other imaging modalities, such as PET and fNIRS. In addition to the article search via PubMed, we manually added seven research articles that the searched articles cited but were not identified through the PubMed search. You can find the full list of the surveyed papers in **Supplementary Table 5**. Fifty-seven articles were included for further analyses.

We categorized the predictive models reported in surveyed articles based on the following aspects: (1) measurement tools (EEG, sMRI, or fMRI), (2) populations (clinical or healthy), and the types of clinical pain, (3) prediction tasks (i.e., classification or regression), (4) targets (e.g., classification of pain vs. no pain, prediction of pain intensity, etc.), (5) model levels (e.g., idiographic model or population-level model), (6) (train and test) data levels (e.g., trial level, run level, etc.), (7) spatial scales (e.g., single region, brain-wide, etc.), (8) experimental tasks (e.g., resting state, phasic pain, etc.), (9) feature types (e.g., activation pattern, connectivity, etc.), (10) algorithms (e.g., linear support vector machine, linear regression, etc.), (11) validation methods (e.g., k-fold cross validation, leave-one out validation, etc.), (12) sample sizes. The full list of the aspects and categories can be found in **Supplementary Table 6**. For the prediction task, we excluded the multiclass classification task because there was only one model for this category (Vijayakumar et al., 2017), and thus all models fell into binary classification or regression. For model performance, we chose to compare the classification accuracy and prediction-outcome correlation (i.e., a correlation between the predicted and actual values), which were the main performance metric that most of the surveyed models adopted (89.5% and 64.3%, respectively; **Supplementary Figure 1**). The final comparisons included the 129 training models and 44 independent tests from 57 studies.

After we divided the model performance into two categories based on prediction tasks (i.e., binary classification or regression), we first compared the performances based on the modeling targets (i.e., what the models are designed to predict). Next, given that most of the models (63.6%) focused on the following target categories, including ‘Pain vs. no pain’ and ‘Low pain vs. high pain’ for binary classification and ‘Pain rating’ for regression, we compared the model performance of these targets for different aspects, including ‘(train) data level’, ‘spatial scales’, ‘model levels’, and ‘sample size.’

### Participants for Benchmarking Analysis

We conducted a benchmarking analysis, in which we compared the performance of models with specific modeling options against those with alternative options to evaluate the influence of modeling choices on performance. We utilized a locally collected, large-scale functional Magnetic Resonance Imaging (fMRI) dataset (*N* = 124), which included painful heat stimuli. Multiple models were trained and tested using this dataset, each incorporating a variety of modeling options. We recruited a total of 137 healthy and right-handed participants with no history of neurological, psychiatric, or chronic pain disorders. Among them, thirteen participants were excluded due to (a) technical issues (e.g., thermal stimulus equipment errors), (b) voluntary discontinuation of the scanning session by participants (e.g., intolerable stimulus), or (c) finding of an abnormal structure in the brain (e.g., Arachnoid cyst). The final number of participants included in the current study was 124 (61 female, age = 22.17 ± 2.69 years (mean ± SD)). This study was approved by the Institutional Review Board at Sungkyunkwan University, and all participants provided written informed consent.

### Experimental Design and Procedure

We conducted the experiment over two days. On the first day, participants visited the laboratory to complete a series of self-report questionnaires. Within 2 weeks, the participants returned to the laboratory, completed another series of self-report questionnaires, and underwent an fMRI experiment. The fMRI experimental procedure consisted of four types of runs: (1) resting-state, (2) thermal stimulus without movie stimuli, (3) thermal stimulus preceded by a short (20-sec) movie clip, and (4) oral capsaicin stimulus. In the current study, we used only the runs involving thermal stimulus, i.e., (2) and (3) above, and the structural scans.

The thermal stimulation runs (with or without movie stimuli) consisted of 12 trials. In each trial, participants experienced a thermal stimulus for 12 seconds (for details, see “Thermal Stimulation” below) while fixating their eye gaze on a cross shown on the screen. After each thermal stimulus, participants rated the magnitude of their painful experience using the generalized Labeled Magnitude Scale (gLMS) (Bartoshuk et al., 2004). This has a 0-1 numerical continuous rating scale with anchors of “Not at all” (0), “A little bit” (0.061), “Moderately” (0.172), “Strongly” (0.354), “Very Strongly” (0.533), and “Most (Strongest imaginable sensation/unpleasantness of any kind)” (1). Participants completed a total of eight runs of thermal stimulation, two of which were the thermal stimulation runs without a movie and six runs with a movie. In the case of the thermal stimulation run with a movie clip, a 20-second movie clip was shown before the thermal stimulation. Figure 6a shows the trial structure with time information. The movie clips were from a Korean film, titled “Summer, Bus” (available at https://youtu.be/-MliIE5PGrI). We split the 12-minute movie into 20-second video clips (i.e., 36 movie clips). The sequence of the experimental runs was structured as follows: the initial run was the thermal stimulation run without a movie, followed by six consecutive runs of thermal stimulation with a movie clip, and concluding with a final run of thermal stimulation without a movie. Each participant received a total of 96 thermal stimuli (i.e., 8 runs and 12 trials for each run). Lastly, for 24 participants, we had to discard one or two runs that had some technical issues (e.g., sound or thermode malfunction, etc.).

### Thermal Stimulation

In the thermal stimulation runs, we delivered thermal stimulation using MRI-compatible PATHWAY ATS (Advance Thermal Stimulation) system (Medoc Ltd., Israel) with a 16×16 mm^2^ thermode. We marked four different sites on the left forearm of each participant for thermal stimulation. For each run, one of the four sites was selected, and the same site was never used in two subsequent runs. The order of the stimulation sites was counterbalanced across participants. We delivered thermal stimulation with fixed temperatures ranging from 45 to 47.5°C in 0.5°C increments. Before the start of each run, we applied the highest temperature (i.e., 47.5°C) on the skin site to avoid the site-specific habituation effects in the middle of runs. On each trial, the stimulation was delivered for 12 seconds (2.5 seconds ramp-up, 7 seconds at plateau temperature, 2.5 seconds ramp-down) from the baseline temperature (32°C).

### fMRI Data Acquisition and Preprocessing

The fMRI data were collected using a 3T Siemens Prisma scanner at the Center for Neuroscience Imaging Research, Institute for Basic Science, Sungkyunkwan University. Structural T1-weighted images were obtained using magnetization-prepared rapid gradient echo sequence (0.7 × 0.7 × 0.7 mm^3^ voxel size, repetition time (TR): 2,400 ms, echo time (TE): 2.34 ms, slice thickness: 0.70 mm, flip angle: 8°, field of view (FoV): 224 × 224 mm^2^, inversion time (TI): 1,150 ms). Functional data were then acquired using gradient echo-planar imaging (EPI) sequence (2.7 × 2.7 × 2.7 mm^3^ voxel size, repetition time (TR): 460 ms, echo time (TE): 27.20 ms, flip angle: 44° slice thickness: 2.7 mm, slices, field of view (FoV): 220 × 220 mm^2^, order of slice accession: interleaved). The first eighteen image volumes of each run were removed prior to image preprocessing for image intensity stabilization. Structural and functional MRI data were preprocessed using our in-house preprocessing pipeline (https://github.com/cocoanlab/humanfmri_preproc_bids) based on Statistical Parametric Mapping 12 (SPM12) software (http://www.fil.ion.ucl.ac.uk/spm/software/spm12), FMRIB Software Library (https://fsl.fmrib.ox.ac.uk/fsl/fslwiki/) and ICA-AROMA (ICA-based strategy for Automatic Removal Of Motion Artifacts) software (https://github.com/maartenmennes/ICA-AROMA). Structural T1-weighted images were co-registered to the functional image for each subject and then normalized to the Montreal Neurological Institute (MNI) space. For functional EPI preprocessing, the pipeline included the following steps: motion correction (realignment), distortion correction using FSL’s topup, co-registration, spatial normalization to Montreal Neurological Institute (MNI) space using structural T1-weighted images with the interpolation to 2×2×2 mm^3^ voxels, spatial smoothing with a Gaussian kernel (5 mm Full-Width Half-Maximum), and independent component analysis (ICA) to automatically detect and remove participant-specific, motion-related artifacts (ICA-AROMA) (Pruim et al., 2015). In a quality control (QC) phase, a few runs were excluded based on the following two criteria based on framewise displacement (FD): (1) the average FD of a run exceeds 0.2 mm, and (2) the FD of any volume was greater than 5 mm in a run (Power et al., 2012; Power et al., 2014).

### Single Trial Analysis

Before conducting the predictive modeling analysis, we estimated single-trial response magnitudes for each voxel using a general linear model (GLM) with separate regressors for each trial, as in the “beta series” approach (Rissman et al., 2004) (Figure 6b). We constructed each trial regressor for movie watching, pain anticipation, and heat stimulation with a boxcar convolved with SPM12’s canonical hemodynamic response function (HRF). We also included one regressor for the pain rating period for each run. In the preprocessing, since we already removed participant-specific, motion-related artifacts in ICA-AROMA, we additionally regressed out only the following nuisance covariates—five principal components of WM and CSF signal and a linear trend. We then calculated trial-by-trial variance inflation factors (VIFs), which measure design-induced uncertainty due to collinearity with nuisance regressors. This step was crucial to identify trials potentially influenced by artifacts. Trials with VIFs exceeding 3 were excluded from further analyses. On average, 0.1371 trials were excluded per participant due to high VIFs, with a standard deviation of 0.7686. The single-trial beta maps served as inputs for predictive modeling.

### Predictive modeling

We trained predictive models for benchmarking analysis based on single-trial beta images. There were two types of tasks: (1) binary classification of high pain vs. low pain, and (2) regression of pain ratings (Figure 6c). For binary classification, we defined “high” pain as heat stimulus levels 5 and 6 and “low” pain as heat stimulus levels 1 and 2. We trained classifier models using linear support vector machines (linear SVMs) implemented in ‘fitcsvm.m’ function from the MATLAB Statistics and Machine Learning Toolbox (https://www.mathworks.com/help/stats/support-vector-machine-classification.html). An SVM classifies data by identifying the optimal hyperplane that separates data points of one class from those of the other class, with distances from the hyperplane indicating the likelihood that an input belongs to one class or the other class (Hastie et al., 2009). For a soft margin parameter, we used the default value of C = 1. For regression, we trained models to predict pain ratings provided by participants using the gLMS. The principal component regression (PCR) algorithm was employed to estimate pain ratings from single-trial fMRI data. The PCR begins by performing principal components analysis (PCA) on the input data to reduce its dimensionality and then linearly fits the component scores to the training data. We selected the minimum number of principal components that explained more than 80% of the variance.

We partitioned the data into training (*n* = 80) and testing datasets (*n* = 44). To guarantee that the training and testing datasets were comparable in terms of the outcome variable’s distribution, we employed a random shuffling for the participants. The final division of datasets was selected based on the criterion of having non-significant differences in pain ratings between the datasets, as determined by two-sample *t*-tests. We did not use any of the testing data during the model training, and all testing results were based on the *n* = 44 hold-out independent test datasets.

### Benchmarking analysis

In the benchmarking analysis, we selected four benchmark scopes that corresponded with the four aspects defined in the survey: (1) Data level, (2) Spatial scale, (3) Model level, and (4) Sample size.

1. **Data level (**Figure 7): The “data level” indicates the number of trials that were averaged for model training and testing. Each participant had a total of 96 single-trial beta estimates and pain ratings if there was no issue, such as technical problems or high VIFs. The 96 trials consisted of two repeats of the same stimulus within a run, a total of 8 runs, and 6 stimulus intensity levels (i.e., 96 = 2×8×6). We established five data levels by averaging different numbers of trials in the training and testing datasets, respectively. First, ‘No Average’ means that we used a total of 96 single-trial beta images and pain ratings without averaging (i.e., trial level). Second, we averaged two trials with a stimulus with the same intensity within a run, resulting in 48 brain images and pain ratings per participant (i.e., run level). Third, we averaged 4 trials with the same intensity across two runs, generating 24 brain images and pain ratings per participant. Fourth, we averaged 8 trials with the same intensity from odd runs (i.e., run 1, 3, 5, 7) and even runs (i.e., run 2, 4, 6, 8), which yielded 12 brain images and pain ratings per participant. Finally, we averaged 16 trials (i.e., two repeats within a run and eight repeats across runs), resulting in 6 brain images and pain ratings per participant (i.e., condition level). In addition, we performed an additional analysis with the matched number of testing data (which was 6 data points) for the different test data levels for fair comparisons across different test data levels. For the comparisons of different data levels, we fixed other benchmark scopes—*gray matter* for the spatial scale, *population-level* modeling for the model level, and *full data* (training data *n* = 80) for sample size.
2. **Spatial scale (**Figure 8): To examine the effects of the spatial scale, we used (1) 21 predefined pain-predictive regions-of-interest (ROIs) obtained from a previous study (Kohoutova et al., 2022) (**Supplementary Figure 2**), (2) a meta-analytic map associated with the term “pain” obtained from Neurosynth.org (association test; downloaded on 1 Nov 2021) (Yarkoni et al., 2011), and (3) a gray matter mask. In the benchmarking analysis for the spatial scale, we tested the impact of increasing spatial scale by randomly selecting one pain-predictive region out of 21 ROIs, then randomly adding more regions incrementally (1, 3, 6, 10, and 15) until all 21 ROIs were included for model training and testing. We repeated this process 100 times and obtained the mean accuracy for each spatial scale and for each participant. For the comparisons of different spatial scales, we fixed other benchmark scopes—*run level* for the data level, *population-level* modeling for the model level, and *full data* (training data *n* = 80) for sample size.
3. **Model level (**Figure 9): We compared idiographic versus population-level predictive models. In this analysis, the sample size of the training dataset was reduced to *n* = 61 because we needed the data to have the complete 8-run data for the analysis described below. For idiographic modeling (i.e., within-individual prediction modeling), we trained a model based on 6-run data and tested the model on the remaining two-run data. For the data split, we also used the two-sample *t*-test to ensure that there was no significant difference in the outcome variable (i.e., pain ratings) between the training and testing datasets. We also trained one population-level model based on 6-run data concatenated across all 61 participants. We tested these two types of models on two different types of testing datasets. One was the remaining two hold-out run data from 61 participants as described above, and the other was all run data from 44 hold-out participants. To test the idiographic models on the 44 hold-out participants’ data, we averaged the idiographic models to construct one predictive model. For the comparisons of the results, we fixed other benchmark scopes—*trial level* for the data level, *gray matter* for the spatial scale, and *n* = 61 for sample size.
4. **Sample size (**Figure 10): Lastly, we evaluated the impact of varying sample sizes using a random selection procedure akin to that employed for evaluating spatial scale. Specifically, we examined the impact of increasing sample sizes by randomly selecting 10 participants from the total pool of 80 participants, then incrementally adding more participants in steps (10, 20, 30, …, 70) until all 80 participants were included in the model training and testing. We repeated this process 100 times and obtained the mean accuracy for each sample size and for each iteration. For the comparisons of different sample sizes, we fixed other benchmark scopes—*run level* for the data level, *population-level* for the model level, and *gray matter* for the spatial scale.

## Supplementary Materials

Supplementary Figures 1-2

Supplementary Tables 1-6

## Author Contributions

S.L. and C.-W.W. designed the experiment. D.H.L. collected the data. D.H.L. and S.L. preprocessed the data. D.H.L. and C.-W.W. analyzed the data, interpreted the results, and wrote the manuscript. C.-W.W. edited the manuscript and provided the supervision.

## Data Availability

All data that were used to generate main figures will be shared upon publication through a Zenodo repository.

## Code Availability

The code for generating the main figures will be shared upon publication through a Zenodo repository. In-house Matlab codes for fMRI data analyses are available at https://github.com/canlab/CanlabCore.

## Acknowledgements

This work was supported by IBS-R015-D1 (Institute for Basic Science; to C.-W.W.) and 2021M3E5D2A01022515 (National Research Foundation of Korea; to C.-W.W.).

