## Supplementary information for "Decoding Pain: Uncovering the Factors that Affect Performance of Neuroimaging-Based Pain Models"

**This PDF file includes:**

1. Supplemental Figures 1-2 (p. 2-3)

2. Supplemental Tables 1-6 (p. 4-13)

3. Supplemental References (p. 14)

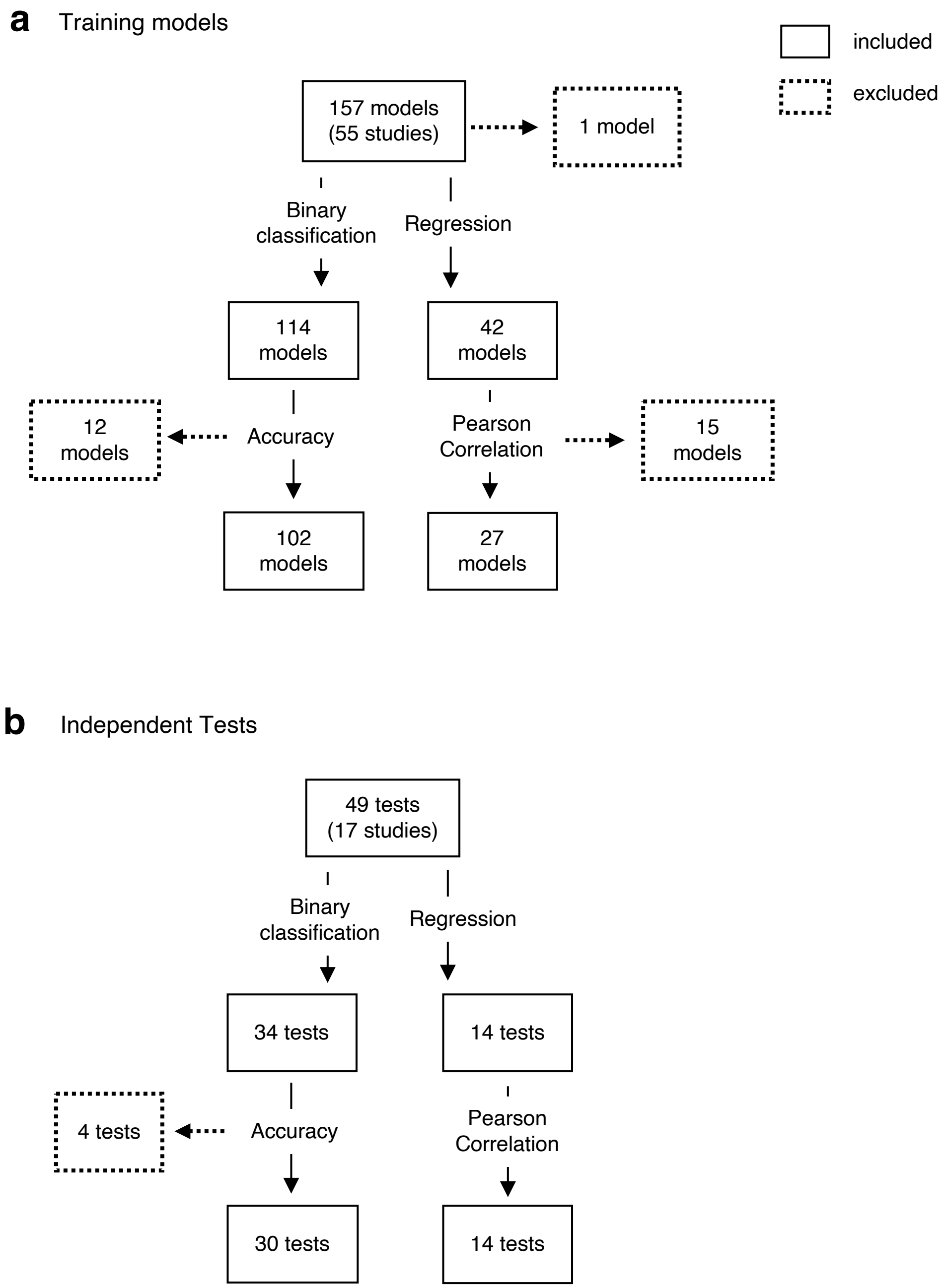


**Supplementary Figure 1. Selection Flowchart for Training Models and Independent Tests.** This flowchart details the criteria and process for the inclusion of (a) training models and (b) independent tests from the literature survey. We included training models and independent tests that reported accuracy using classification accuracy or Pearson correlations. The boxes with solid lines indicate the number of test performances included in the survey. The boxes with dotted lines indicate the number of test performances excluded from the survey, which included measures such as sensitivity or mean squared error. Also, we excluded one multi-class classification task from the training model

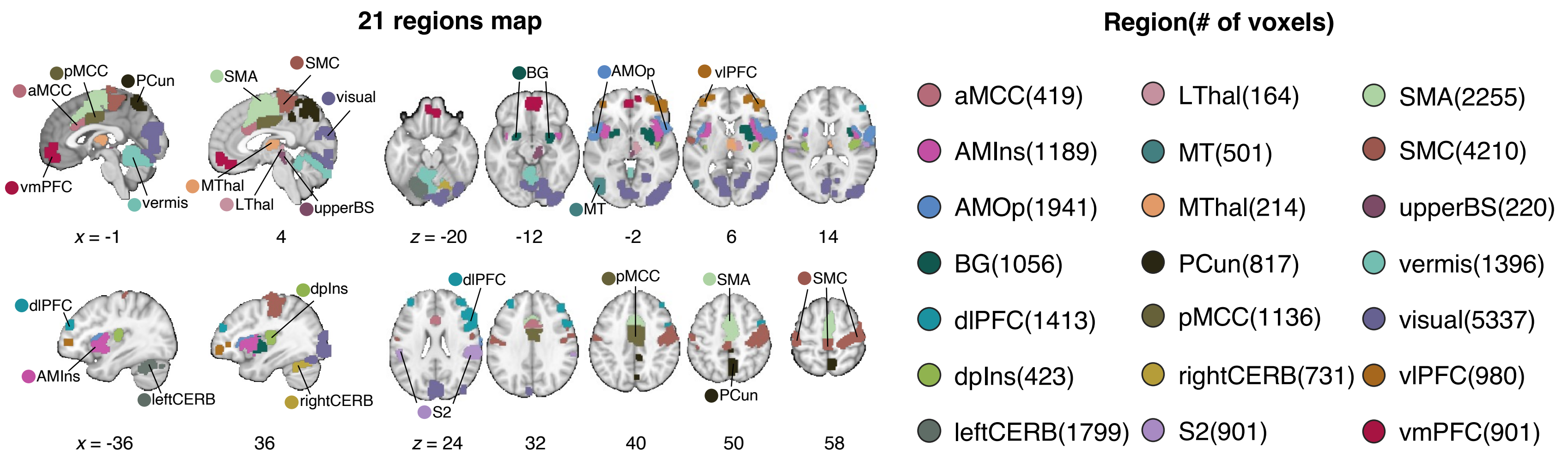


**Supplementary Figure 2. Pre-defined 21 pain-predictive regions**. We used pre-defined regions-of-interest (ROIs) that were identified from a previous study (Kohoutova et al., 2022). On the left, colored areas illustrate the spatial extent of the brain regions. Each region included a different number of voxels, which is provided on the right. aMCC, anterior midcingulate cortex; AMIns, anterior middle insula; AMOp, anterior middle operculum; BG, basal ganglia; dlPFC, dorsal lateral prefrontal cortex; dpIns, dorsal posterior insula; leftCERB, left cerebellum; LThal, lateral thalamus; MT, middle temporal area; MThal, middle thalamus; PCun, precuneus; pMCC, posterior midcingulate cortex; rightCERB, right cerebellum; S2, secondary somatosensory cortex; SMA, supplementary motor area; SMC, sensorimotor cortex; upperBS, upper brainstem; visual, visual cortex; vlPFC, ventrolateral prefrontal cortex; vmPFC, ventromedial prefrontal cortex.

**Supplementary Table 1. Model performance for data levels**

| Benchmark | Train data level | Test data level | Binary Classification | | | | | | Regression | | | |
|---|---|---|---|---|---|---|---|---|---|---|---|---|
| | | | Accuracy | Accuracy multi-level GLM | Sensitivity | Specificity | AUC | PPV | *r* | *r* multi-level GLM | RMSE | $R^2$ |
| Train data level | no avg | no avg | 85.05 ±1.31 | beta = -1.09 $z$ = -3.53 $p$ = 0.00042 two-tailed, bootstrap test | 87.53 ±2.52 | 82.56 ±2.44 | 95.44 ±0.84 | 85.72 ±1.61 | 0.6898 ±0.0117 | beta = -0.03 $z$ = -3.75 $p$ = 0.00018 two-tailed, bootstrap test | 0.1711 ±0.0070 | -0.2985 ±0.1565 |
| | 2 | | 84.53 ±1.32 | | 86.58 ±2.60 | 82.47 ±2.37 | 95.45 ±0.75 | 85.16 ±1.48 | 0.6782 ±0.0178 | | 0.1768 ±0.0073 | -0.3907 ±0.1616 |
| | 4 | | 83.17 ±1.43 | | 85.82 ±2.86 | 80.50 ±2.62 | 95.04 ±0.86 | 83.88 ±1.58 | 0.6618 ±0.0180 | | 0.1868 ±0.0077 | -0.5845 ±0.1900 |
| | 8 | | 81.70 ±1.44 | | 85.73 ±2.79 | 77.64 ±3.06 | 94.23 ±0.88 | 82.45 ±1.74 | 0.6232 ±0.0197 | | 0.2071 ±0.0094 | -1.0013 ±0.2646 |
| | 16 | | 80.31 ±1.45 | | 84.58 ±2.94 | 76.03 ±2.93 | 92.91 ±0.85 | 80.44 ±1.61 | 0.5784 ±0.0231 | | 0.2014 ±0.0080 | -0.8452 ±0.2270 |
| | no avg | 16 | 93.75 ±2.17 | beta = -5.09 $z$ = -4.50 $p$ = 0.00001 two-tailed, bootstrap test | 95.45 ±3.18 | 92.05 ±3.23 | 100.00 ±0.00 | 94.84 ±2.01 | 0.9737 ±0.0036 | beta = -0.003 $z$ = -3.58 $p$ = 0.00034 two-tailed, bootstrap test | 0.1233 ±0.0087 | -0.3647 ±0.2499 |
| | 2 | | 93.75 ±2.17 | | 94.32 ±3.34 | 93.18 ±3.08 | 100.00 ±0.00 | 95.63 ±1.89 | 0.9746 ±0.0031 | | 0.1243 ±0.0091 | -0.4015 ±0.2460 |
| | 4 | | 93.18 ±2.35 | | 92.05 ±3.96 | 94.32 ±2.92 | 100.00 ±0.00 | 96.34 ±1.80 | 0.9747 ±0.0033 | | 0.1317 ±0.0096 | -0.6263 ±0.2892 |
| | 8 | | 90.34 ±2.72 | | 92.05 ±3.96 | 88.64 ±4.26 | 100.00 ±0.00 | 93.09 ±2.47 | 0.9698 ±0.0043 | | 0.1420 ±0.0119 | -1.0094 ±0.3770 |
| | 16 | | 89.77 ±2.73 | | 89.77 ±4.17 | 89.77 ±4.17 | 100.00 ±0.00 | 93.90 ±2.38 | 0.9509 ±0.0092 | | 0.1337 ±0.0102 | -0.6608 ±0.2980 |

(Supplementary Table 1 continues on the next page)

| Benchmark | Train data level | Test data level | Binary Classification | | | | | | | Regression | | |
|---|---|---|---|---|---|---|---|---|---|---|---|---|
| | | | Accuracy | Accuracy multi-level GLM | Sensitivity | Specificity | AUC | PPV | $r$ | $r$ multi-level GLM | RMSE | $R^2$ |
| Test data level | no avg | no avg | 85.05 ±1.31 | beta = 2.35 $z$ = 3.43 $p$ = 0.00061 two-tailed, bootstrap test | 87.53 ±2.52 | 82.56 ±2.44 | 95.44 ±0.84 | 85.72 ±1.61 | 0.6898 ±0.0117 | beta = 0.07 $z$ = 4.43 $p$ = 0.00001 two-tailed, bootstrap test | 0.1711 ±0.0070 | -0.2985 ±0.1565 |
| | | 2 | 88.36 ±1.52 | | 90.38 ±2.35 | 86.33 ±2.75 | 97.78 ±0.64 | 89.41 ±1.82 | 0.7542 ±0.0175 | | 0.1571 ±0.0073 | -0.3256 ±0.1866 |
| | | 4 | 91.57 ±1.62 | | 92.33 ±2.70 | 90.81 ±2.49 | 99.27 ±0.51 | 93.00 ±1.71 | 0.7903 ±0.0184 | | 0.1473 ±0.0077 | -0.4267 ±0.2173 |
| | | 8 | 92.90 ±2.05 | | 94.89 ±2.88 | 90.91 ±3.26 | 100.00 ±0.00 | 93.96 ±1.97 | 0.9235 ±0.0093 | | 0.1291 ±0.0082 | -0.3600 ±0.2354 |
| | | 16 | 93.75 ±2.17 | | 95.45 ±3.18 | 92.05 ±3.23 | 100.00 ±0.00 | 94.84 ±2.01 | 0.9737 ±0.0036 | | 0.1233 ±0.0087 | -0.3647 ±0.2499 |
| | 16 | no avg | 80.31 ±1.45 | beta = 2.28 $z$ = 3.40 $p$ = 0.00068 two-tailed, bootstrap test | 84.58 ±2.94 | 76.03 ±2.93 | 92.91 ±0.85 | 80.43 ±1.60 | 0.5784 ±0.0231 | beta = 0.09 $z$ = 3.93 $p$ = 0.00009 two-tailed, bootstrap test | 0.2013 ±0.0079 | -0.8452 ±0.2269 |
| | | 2 | 84.29 ±1.65 | | 87.66 ±2.83 | 80.92 ±3.23 | 96.06 ±0.75 | 85.43 ±1.86 | 0.6633 ±0.0240 | | 0.1802 ±0.0084 | -0.7834 ±0.2498 |
| | | 4 | 86.74 ±2.13 | | 89.20 ±3.46 | 84.28 ±3.66 | 98.88 ±0.52 | 89.16 ±2.15 | 0.7334 ±0.0239 | | 0.1629 ±0.0090 | -0.8009 ±0.2823 |
| | | 8 | 88.64 ±2.63 | | 90.91 ±3.82 | 86.36 ±4.34 | 99.93 ±0.07 | 91.44 ±2.46 | 0.8720 ±0.0202 | | 0.1416 ±0.0097 | -0.6915 ±0.2901 |
| | | 16 | 89.77 ±2.73 | | 89.77 ±4.17 | 89.77 ±4.17 | 100 ±0.00 | 93.90 ±2.38 | 0.9509 ±0.0092 | | 0.1336 ±0.010 | -0.6608 ±0.2979 |

(Supplementary Table 1 continues on the next page)

| Benchmark | Train data level | Test data level | Binary Classification | | | | | | Regression | | | |
|---|---|---|---|---|---|---|---|---|---|---|---|---|
| | | | Accuracy | Accuracy multi-level GLM | Sensitivity | Specificity | AUC | PPV | $r$ | $r$ multi-level GLM | RMSE | $R^2$ |
| Test data level (with matched # data across test data levels) | no avg | no avg | 87.50 ±2.37 | beta = 4.56 $z$ = 3.79 $p$ = 0.00015 two-tailed, bootstrap test | 92.04 ±3.22 | 82.95 ±4.28 | 98.86 ±0.54 | 89.14 ±2.59 | 0.7806 ±0.0276 | beta = 0.04 $z$ = 4.65 $p$ = 0.000003 two-tailed, bootstrap test | 0.1589 ±0.0081 | -1.0787 ±0.4316 |
| | | 2 | 88.64 ±2.50 | | 95.45 ±3.17 | 81.81 ±4.32 | 100.00 ±0.00 | 88.09 ±2.68 | 0.8522 ±0.0204 | | 0.1553 ±0.0090 | -0.9456 ±0.4052 |
| | | 4 | 92.61 ±2.08 | | 92.04 ±3.61 | 93.18 ±2.61 | 100.00 ±0.00 | 95.23 ±1.82 | 0.8871 ±0.0181 | | 0.1479 ±0.0110 | -0.7232 ±0.4057 |
| | | 8 | 93.18 ±2.05 | | 94.31 ±2.91 | 92.04 ±3.22 | 100.00 ±0.00 | 94.96 ±1.96 | 0.9556 ±0.0061 | | 0.1261 ±0.0081 | -0.4056 ±0.3440 |
| | | 16 | 93.75 ±2.17 | | 95.45 ±3.17 | 92.04 ±3.22 | 100.00 ±0.00 | 94.84 ±2.00 | 0.9737 ±0.0036 | | 0.1233 ±0.0087 | -0.3647 ±0.2499 |
| | 16 | no avg | 81.25 ±2.94 | beta = 3.49 $z$ = 3.18 $p$ = 0.00147 two-tailed, bootstrap test | 85.23 ±4.48 | 77.27 ±5.00 | 96.59 ±1.31 | 84.55 ±3.16 | 0.6498 ±0.0528 | beta = 0.06 $z$ = 3.82 $p$ = 0.00013 two-tailed, bootstrap test | 0.1887 ±0.0102 | -1.9572 ±0.5072 |
| | | 2 | 84.09 ±3.05 | | 87.50 ±4.63 | 80.68 ±5.20 | 99.15 ±0.48 | 87.92 ±3.04 | 0.7995 ±0.0266 | | 0.1823 ±0.0116 | -1.6389 ±0.4568 |
| | | 4 | 85.80 ±2.86 | | 87.50 ±4.63 | 84.09 ±4.53 | 100.00 ±0.00 | 89.58 ±2.78 | 0.8650 ±0.0173 | | 0.1595 ±0.0125 | -0.9096 ±0.3439 |
| | | 8 | 86.93 ±2.99 | | 89.77 ±4.17 | 84.09 ±5.08 | 100.00 ±0.00 | 90.65 ±2.85 | 0.9240 ±0.0100 | | 0.1410 ±0.0102 | -0.7950 ±0.4213 |
| | | 16 | 89.77 ±2.73 | | 89.77 ±4.17 | 89.77 ±4.17 | 100.00 ±0.00 | 93.90 ±2.38 | 0.9509 ±0.0092 | | 0.1337 ±0.0102 | -0.6608 ±0.2980 |

*Note.* This table, related to Figure 7, displays performance metrics for different data levels, including the mean and standard error of the mean (s.e.m.). AUC, Area under the ROC curve; PPV, Positive Predictive Value; $r$, Pearson correlation between the actual and predicted values; RMSE, Root Mean Square Error; $R^2$, r-squared (or explained variance)

**Supplementary Table 2. Model performance for spatial scales**

| Spatial Scale | | Binary Classification | | | | | | | Regression | | |
|---|---|---|---|---|---|---|---|---|---|---|---|
| | | Accuracy | Accuracy multi-level GLM<br>Paired *t*-test | Sensitivity | Specificity | AUC | PPV | *r* | *r* multi-level GLM<br>Paired *t*-test | RMSE | $R^2$ |
| Number of Regions | 1 | 68.79 ±4.92 | | 67.44 ±13.42 | 70.13 ±10.90 | 80.58 ±5.53 | 71.54 ±5.37 | 0.4583 ±0.1048 | | 0.177 ±0.0468 | -0.5132 ±1.0586 |
| | 3 | 73.01 ±5.15 | | 73.33 ±14.74 | 72.69 ±12.69 | 86.22 ±5.12 | 75.44 ±6.71 | 0.5658 ±0.1057 | | 0.1737 ±0.0460 | -0.5239 ±1.2164 |
| | 6 | 78.67 ±6.46 | beta = 4.25<br>*z* = 3.66<br>*p* = 0.00025<br>two-tailed,<br>bootstrap test | 80.73 ±15.70 | 76.61 ±14.44 | 92.17 ±4.54 | 80.46 ±8.30 | 0.6242 ±0.1063 | beta = 0.06<br>*z* = 3.58<br>*p* = 0.00034<br>two-tailed,<br>bootstrap test | 0.1717 ±0.0480 | -0.5354 ±1.3468 |
| | 10 | 82.56 ±7.51 | | 85.57 ±15.09 | 79.54 ±15.63 | 95.36 ±3.84 | 83.61 ±9.48 | 0.6663 ±0.1084 | | 0.1707 ±0.0504 | -0.5403 ±1.3986 |
| | 15 | 85.27 ±8.14 | | 89.03 ±14.35 | 81.52 ±16.35 | 96.93 ±3.10 | 85.58 ±10.28 | 0.6973 ±0.1103 | | 0.1694 ±0.0514 | -0.5251 ±1.3922 |
| Brain-wide masks | 21 | 87.31 ±9.32 | | 91.59 ±13.76 | 83.03 ±18.21 | 97.6 ±2.66 | 86.84 ±11.89 | 0.7105 ±0.1231 | | 0.1685 ±0.0534 | -0.5081 ±1.4036 |
| | NP | 86.96 ±8.92 | **21 vs. NP:**<br>$t(43) = 0.2882$,<br>$p = 0.7746$<br>**GM vs. NP:**<br>$t(43) = 0.9677$,<br>$p = 0.3386$<br>**GM vs. 21:**<br>$t(43) = 0.9808$,<br>$p = 0.3322$ | 89.36 ±13.68 | 84.57 ±16.74 | 97.38 ±2.59 | 87.15 ±10.46 | 0.7227 ±0.0971 | **21 vs. NP:**<br>$t(43) = -0.7642$,<br>$p = 0.4489$<br>**GM vs. NP:**<br>$t(43) = -1.9625$,<br>$p = 0.0562$<br>**GM vs. 21:**<br>$t(43) = -2.6264$,<br>$p = 0.0119$ | 0.1633 ±0.0553 | -0.4317 ±1.3331 |
| | GM | 88.27 ±10.46 | | 89.51 ±17.79 | 87.03 ±18.06 | 97.96 ±3.27 | 89.98 ±11.68 | 0.7455 ±0.1196 | | 0.1613 ±0.0503 | -0.4008 ±1.2535 |

*Note.* This table, related to Figure 8, displays performance metrics for different spatial scales, including the mean and standard deviation. AUC, Area under the ROC curve; PPV, Positive Predictive Value; *r*, Pearson correlation between the actual and predicted values; RMSE, Root Mean Square Error; $R^2$, r-squared (or explained variance)

**Supplementary Table 3. Model performance for model levels**

| Train | Test | Binary Classification | | | | | | Regression | | | |
|---|---|---|---|---|---|---|---|---|---|---|---|
| | | Accuracy | Accuracy Paired *t*-test | Sensitivity | Specificity | AUC | PPV | *r* | *r* Paired *t*-test | RMSE | $R^2$ |
| Idiographic model | Within-individual run-level test | 84.43 ±11.27 | **Id vs. Po**: *t*(120) = 0.2710, *p* = 0.7868 | 82.99 ±14.80 | 85.86 ±16.84 | 94.56 ±7.00 | 87.21 ±13.96 | 0.6448 ±0.1844 | **Id vs. Po**: *t*(120) = -1.2872, *p* = 0.2005 | 0.1241 ±0.0403 | 0.3592 ±0.2615 |
| Population-level model | | 83.81 ±13.66 | | 79.71 ±25.02 | 87.91 ±18.54 | 94.43 ±10.86 | 88.24 ±17.63 | 0.6875 ±0.1819 | | 0.1569 ±0.0577 | -0.2891 ±1.5668 |
| Average of idiographic models | Independent test | 86.76 ±6.30 | **Avg Id vs. Po**: *t*(86) = 1.1938, *p* = 0.2358 | 85.79 ±10.08 | 87.74 ±8.07 | 92.08 ±5.47 | 87.96 ±7.01 | 0.6051 ±0.1252 | **Avg Id vs. Po**: *t*(86) = -2.8129, *p* = 0.0061 | 0.1832 ±0.0661 | -0.3567 ±1.0632 |
| Population-level model | | 84.79 ±8.98 | | 86.41 ±17.64 | 83.16 ±16.49 | 95.67 ±4.89 | 86.02 ±10.39 | 0.6807 ±0.1271 | | 0.1732 ±0.0514 | -0.3334 ±1.1137 |

*Note.* This table, related to Figure 9, displays performance metrics for different model levels, including the mean and standard deviation. AUC, Area under the ROC curve; PPV, Positive Predictive Value; *r*, Pearson correlation between the actual and predicted values; RMSE, Root Mean Square Error; $R^2$, r-squared (or explained variance)

**Supplementary Table 4. Model performance for sample sizes**

| Sample Size | Binary Classification | | | | | | Regression | | | |
|---|---|---|---|---|---|---|---|---|---|---|
| | Accuracy | Accuracy multi-level GLM | Sensitivity | Specificity | AUC | PPV | *r* | *r* multi-level GLM | RMSE | $R^2$ |
| 10 | 83.50 ±0.58 | | 81.12 ±3.44 | 85.88 ±2.67 | 96.18 ±0.70 | 87.88 ±1.59 | 0.6981 ±0.0164 | | 0.1664 ±0.0051 | -0.5135 ±0.1346 |
| 20 | 85.39 ±1.09 | | 84.48 ±2.40 | 86.31 ±2.61 | 97.24 ±0.28 | 89.09 ±1.86 | 0.7266 ±0.0127 | | 0.1638 ±0.0052 | -0.4898 ±0.1333 |
| 30 | 85.94 ±0.83 | beta = 0.68 $z = 3.96$ $p = 0.00008$ two-tailed, bootstrap test | 85.84 ±1.95 | 86.04 ±2.79 | 97.49 ±0.28 | 88.54 ±2.21 | 0.7376 ±0.0099 | beta = 0.01 $z = 4.02$ $p = 0.00006$ two-tailed, bootstrap test | 0.1626 ±0.0046 | -0.4640 ±0.1114 |
| 40 | 86.86 ±0.70 | | 88.27 ±1.42 | 85.46 ±2.01 | 97.8 ±0.25 | 88.83 ±1.49 | 0.7412 ±0.0089 | | 0.1621 ±0.0038 | -0.4476 ±0.0887 |
| 50 | 87.15 ±0.40 | | 88.39 ±1.43 | 85.91 ±2.03 | 97.88 ±0.22 | 89.09 ±1.61 | 0.7435 ±0.0073 | | 0.1624 ±0.0039 | -0.4452 ±0.0832 |
| 60 | 87.59 ±0.34 | | 88.66 ±1.01 | 86.52 ±1.23 | 97.94 ±0.14 | 89.56 ±0.98 | 0.7449 ±0.0056 | | 0.162 ±0.0028 | -0.4306 ±0.0604 |
| 70 | 87.99 ±0.40 | | 88.97 ±0.54 | 87.01 ±0.85 | 98.01 ±0.12 | 89.87 ±0.57 | 0.7455 ±0.0041 | | 0.1615 ±0.0021 | -0.4129 ±0.0444 |
| 80 | 88.27 | - | 89.51 | 87.03 | 97.96 | 89.98 | 0.7455 | - | 0.1613 | -0.4008 |

*Note.* This table, related to Figure 10, displays performance metrics for different sample sizes, including the mean and standard deviation. AUC, Area under the ROC curve; PPV, Positive Predictive Value; *r*, Pearson correlation between the actual and predicted values; RMSE, Root Mean Square Error; $R^2$, r-squared (or explained variance)

**Supplementary Table 5. The list of research articles included in the literature survey (*N* = 57)**

| Reference | Measurement tool | Population | Clinical pain type | # of training models | # of independent tests |
|---|---|---|---|---|---|
| {Hunter, 2009} | EEG | Clinical | Fibromyalgia(FM) | 2 | - |
| (Marquand et al., 2010) | fMRI | Healthy | - | 12 | - |
| (Prato et al., 2011) | fMRI | Healthy | - | 9 | - |
| (Brown et al., 2011) | fMRI | Healthy | - | 7 | 1 |
| (Brodersen et al., 2012) | fMRI | Healthy | - | 6 | - |
| (Schulz et al., 2012) | EEG | Healthy | - | 5 | - |
| (Graversen et al., 2012) | EEG | Clinical | Chronic pancreatitis | 1 | - |
| (Wager et al., 2013) | fMRI | Healthy | - | 1 | 11 |
| (Bagarinao et al., 2014) | sMRI | Clinical | Chronic Pelvic Pain | 2 | - |
| (Callan, 2014) | fMRI | Clinical | chronic low back pain(cLBP) | 1 | - |
| (Labus et al., 2015) | sMRI | Clinical | Irritable Bowel Syndrome(IBS) | 1 | - |
| (Gram et al., 2015) | EEG | Healthy | - | 2 | - |
| (Robinson et al., 2015) | sMRI | Clinical | Fibromyalgia(FM) | 2 | - |
| (Tetreault et al., 2016) | fMRI | Clinical | osteoarthritis pain(OA) | 1 | 2 |
| (Tu et al., 2016) | EEG+fMRI | Healthy | - | 8 | - |
| (Harte et al., 2016) | fMRI | Clinical | Fibromyalgia(FM) | 2 | 1 |
| (Harper et al., 2016) | fMRI | Clinical | temporomandibular disorders(TMD) | 4 | - |
| (Lopez-Sola et al., 2017) | fMRI | Clinical | Fibromyalgia(FM) | 2 | 5 |
| (Chong et al., 2017) | fMRI | Clinical | Migraine | 1 | - |
| (Vijayakumar et al., 2017) | EEG | Healthy | - | 2 | - |
| (Lopez-Sola et al., 2020) | fMRI | Healthy | - | - | 1 |

(Supplementary Table 5 continues on the next page)

| Reference | Measurement tool | Population | Clinical pain type | # of training models | # of independent tests |
|---|---|---|---|---|---|
| {Lindquist, 2017} | fMRI | Healthy | - | 4 | - |
| (Woo et al., 2017) | fMRI | Healthy | - | 1 | 2 |
| (Misra et al., 2017) | EEG | Healthy | - | 2 | - |
| (Gram et al., 2017) | EEG | Clinical | - | 1 | - |
| (Vuckovic et al., 2018) | EEG | Clinical | Spinal cord injured | 6 | - |
| (Lin et al., 2018) | fMRI | Healthy | - | 1 | - |
| (Zhong et al., 2018) | sMRI | Clinical | Trigeminal Neuralgia | 1 | - |
| (Mano et al., 2018) | fMRI | Clinical | chronic low back pain(cLBP) | 4 | 2 |
| (Okolo & Omurtag, 2018) | EEG | Healthy | - | 3 | - |
| (Furman et al., 2018) | EEG | Healthy | - | 1 | - |
| (Li et al., 2018) | EEG | Healthy | - | 2 | - |
| (Shen et al., 2019) | fMRI | Clinical | chronic low back pain(cLBP) | 1 | 1 |
| (Lee et al., 2019) | fMRI | Clinical | chronic low back pain(cLBP) | 1 | - |
| (Santana et al., 2019) | fMRI | Clinical | Fibromyalgia(FM), back pain | 4 | - |
| (Rogachov et al., 2019) | fMRI | Clinical | Neuropathic pain | 1 | - |
| (Paul et al., 2019) | EEG | Clinical | Fibromyalgia(FM) | 2 | - |
| (Tu et al., 2019) | fMRI | Clinical | chronic low back pain(cLBP) | 2 | - |
| (Jung et al., 2019) | fMRI | Healthy | - | 6 | - |
| (Liu et al., 2019) | fMRI | Clinical | Knee osteoarthritis(KOA) | 1 | - |
| (Chen et al., 2019) | sMRI | Clinical | Primary Dysmenorrhea(PDM) | 2 | - |
| (Zeng et al., 2019) | sMRI | Clinical | Herpes Zoster(HZ) | 1 | - |

(Supplementary Table 5 continues on the next page)

| Reference | Measurement tool | Population | Clinical pain type | # of training models | # of independent tests |
|---|---|---|---|---|---|
| (Brown et al., 2019) | EEG | Healthy | - | 2 | - |
| (Spisak et al., 2020) | fMRI | Healthy | - | 1 | 2 |
| (Frid et al., 2020) | EEG | Clinical | Migraine | 1 | - |
| (Mao et al., 2020) | fMRI | Clinical | Irritable Bowel Syndrome(IBS) | 4 | 4 |
| (Quan et al., 2021) | fMRI | Clinical | Primary Dysmenorrhea(PDM) | 1 | - |
| (Makary et al., 2020) | fMRI | Clinical | chronic low back pain(cLBP) | 1 | 2 |
| (Wei et al., 2022) | EEG | Clinical | Herpes Zoster(HZ) | 4 | - |
| (Geuter et al., 2020) | fMRI | Healthy | - | - | 3 |
| (Huang et al., 2013) | EEG | Healthy | - | 4 | - |
| (Wager et al., 2011) | fMRI | Healthy | - | 5 | - |
| (Liang et al., 2013) | fMRI | Healthy | - | 9 | - |
| (Ung et al., 2014) | sMRI | Clinical | chronic low back pain(cLBP) | 1 | - |
| (Baliki et al., 2012) | fMRI | Clinical | subacute back pain | 1 | 1 |
| (Liang et al., 2019) | fMRI | Healthy | - | 4 | 2 |
| (Lee et al., 2021) | fMRI | Healthy | - | 1 | 8 |

*Note.* EEG, Electroencephalography; fMRI, functional Magnetic Resonance Imaging; sMRI, structural Magnetic Resonance Imaging

**Supplementary Table 6. Categories for the literature survey**

| Aspect | Categories |
| --- | --- |
| Measurement tool | EEG / sMRI / fMRI |
| Population | Clinical / Healthy |
| Prediction task | Binary classification / Multiclass classification / Regression |
| Target | Pain vs. no pain / Pain intensity / Pain sensitivity / Pain patients vs. controls / Pain persistence / Pain severity / Pain patient type / Treatment responder / Treatment effect / Pain site / Non-pain conditions |
| Model level | Idiographic model / Population-level model / Combination |
| Data level | TR level / TR-bin level / Trial level / Run level / Condition level / Individual level |
| Spatial scale | Single voxel / Single region / Combination of regions / Brain wide |
| Experimental task | Resting state / Phasic pain / Tonic pain / Structure |
| Feature type | Activation pattern / Activation mean / Connectivity pattern / Event Related Potentials / Time Frequency / Structural information |
| Algorithm | ANN / CNN / CVAE / DQDA / GPC / GPR / Decision tree / K-NN / LASSO-PCR / LDA / Linear Regression / Linear SVM / Linear SVR / Logistic regression / PLSR / Random Forest / RVM / RVR / SMO-SVM / the v-method / TPOT |
| Validation method | Holdout validation / K-fold CV / Leave-one-trial-out CV / Leave-one-run-out CV / Leave-one-participant-out CV / Leave-two-participant-out CV / Leave-three-participant-out CV |
| Sample size | Number of participants |

*Note.* In the literature survey, we categorized models based on the listed categories. All models correspond to one category for each aspect. EEG, Electroencephalography; fMRI, functional Magnetic Resonance Imaging; sMRI, structural Magnetic Resonance Imaging; ANN, Artificial Neural Networks; CNN, Convolutional Neural Networks; CVAE, Conditional Variational Autoencoder; DQDA, Diagonal Quadratic Discriminate Analysis; GPC, Gaussian process classifier; GPR, Gaussian Process Regression; KNN, K-Nearest Neighbor; LASSO-PCR, the Least Absolute Shrinkage and Selection Operator Principal Component Regression; LDA, Linear Discriminant Analysis; SVM, Support vector machine; SVR, Support vector regression; PLSR, Partial Least Squares Regression; RVM, Relevance Vector Machine; RVR, Relevance Vector Regression; SMO-SVM, Sequential Minimum Optimization Support Vector Machine; CV, Cross-Validation
